# CHAMP1 complex directs heterochromatin assembly and promotes homology-directed DNA repair

**DOI:** 10.1101/2024.09.23.614480

**Authors:** Feng Li, Tianpeng Zhang, Aleem Syed, Amira Elbakry, Noella Holmer, Huy Nguyen, Sirisha Mukkavalli, Roger A. Greenberg, Alan D. D’Andrea

## Abstract

The CHAMP1 complex, a little-known but highly conserved protein complex consisting of CHAMP1, POGZ, and HP1α, is enriched in heterochromatin though its cellular function in these regions of the genome remain unknown. Here we show that the CHAMP complex promotes heterochromatin assembly at multiple chromosomal sites, including centromeres and telomeres, and promotes homology-directed repair (HDR) of DNA double strand breaks (DSBs) in these regions. The CHAMP1 complex is also required for heterochromatin assembly and DSB repair in highly-specialized chromosomal regions, such as the highly-compacted telomeres of ALT (Alternative Lengthening of Telomeres) positive tumor cells. Moreover, the CHAMP1 complex binds and recruits the writer methyltransferase SETDB1 to heterochromatin regions of the genome and is required for efficient DSB repair at these sites. Importantly, peripheral blood lymphocytes from individuals with CHAMP1 syndrome, an inherited neurologic disorder resulting from heterozygous mutations in *CHAMP1*, also exhibit defective heterochromatin clustering and defective repair of local DSBs, suggesting that a defect in DNA repair underlies this syndrome. Taken together, the CHAMP1 complex has a novel role in heterochromatin assembly and the enhancement of HDR in heterochromatin.

## INTRODUCTION

DNA double strand breaks (DSBs), resulting from intrinsic DNA replication errors or from external exposure to DNA damaging agents, can occur at any site throughout the genome. Homology-directed repair (HDR), including classical homologous recombination (HR) and break-induced replication (BIR), functions to repair these DSBs. Some regions of chromosomes may require specific mechanisms of DSB repair. For example, heterochromatic regions of chromosomes, such as centromeres and telomeres, are highly-compacted structures and pose a significant barrier to HDR-mediated DSB repair. Accordingly, cells may require distinct mechanisms to mediate HDR in these chromosomal regions^1–4^.

Heterochromatin regions of the genome are characterized by increased trimethylation of histone H3 on lysine 9 (H3K9me3), a key modification that interacts with histone reader proteins^5^ and is critical for heterochromatin clustering and function^6^. Heterochromatin assembly is regulated by the coordinated activity of writer H3K9 methyltransferases and eraser H3K9 demethylases. These enzymes regulate the level of H3K9me3 in heterochromatin^6,7^, and these mechanisms are coupled to the HDR repair process^8–12^.

Some heterochromatin structures have a unique nucleosome organization, and these structures may also require specialized mechanisms for heterochromatin assembly and local HDR repair. For example, approximately 10-15% of cancers employ a telomerase-independent mechanism known as the alternative lengthening of telomeres (ALT pathway). This process relies on HDR for template driven telomere lengthening by a break induced replication-like mechanism^13,14^. Moreover, ALT tumor cells have a high level of H3K9me3 and a specialized heterochromatic microenvironment that facilitates DNA synthesis and telomere expansion^15,16^. Whether the CHAMP1 complex plays a specific role in ALT telomere maintenance or ALT telomere DSB repair remains unknown.

A previous report^17^ identified a distinct heterochromatin complex-namely, the CHAMP1/POGZ/HP1α complex-with H3K9me3-binding activity. CHAMP1 (CHromosome Alignment Maintaining Phosphoprotein 1), a highly-conserved zinc finger protein, functions as a regulator of chromosome segregation in mitosis and is required for correct alignment of chromosomes on the metaphase plate^18^. The CHAMP1 binding partner POGZ (POGO transposable element with ZNF domain), also a zinc-finger protein, functions in mitotic cell cycle progression and kinetochore assembly^19^. HP1α, a member of the HP1 family of “hub proteins”, interacts with several chromosomal proteins and binds to H3K9me3 through its chromodomain (CD). Whether this CHAMP1/POGZ/HP1α complex regulates heterochromatin assembly or mediates HDR in condensed regions of chromosomes remains unknown.

Humans with *de novo* heterozygous mutations in *CHAMP1*, known as CHAMP1 syndrome, present with a neurodevelopmental disorder characterized by intellectual disability, behavioral symptoms, and distinct dysmorphic features ^20–24^. Interestingly*, de novo* heterozygous mutations in the *POGZ* gene also result in a rare but highly-related neurodevelopment syndrome called the White Sutton Syndrome (WSS) ^25–33^. Whether these inherited mutations result from defects in heterochromatin assembly or HDR in heterochromatin remains unknown.

Recent studies have elucidated at least one mechanism by which CHAMP1 and POGZ promote HR ^34–36^. CHAMP1 has a unique, high affinity site for the DNA repair HORMA protein, REV7. By binding to REV7, the CHAMP1/POGZ complex steals REV7 from the Shieldin (REV7/SHLD) complex^36^. The Shieldin complex normally blocks DNA end resection, promotes NHEJ repair, and inhibits HR repair ^37–41^. By binding to REV7, the CHAMP1/POGZ complex increases DSB end resection and promotes HR repair. Whether this mechanism of enhanced HR repair occurs in heterochromatin, or whether its use is confined to euchromatic regions, remains unknown.

In the current study we show that the CHAMP1/POGZ/HP1α heterochromatin complex performs a direct role in heterochromatin clustering and HDR repair. The complex promotes the deposition of H3K9me3, the clustering of heterochromatin, and the recruitment of HDR proteins (both HR proteins and BIR proteins) to DSBs at heterochromatic sites. While the complex subunits promote heterochromatin clustering at multiple sites, including centromeres and telomeres, it has a distinct and obligate role in maintaining ALT telomeres. Depletion of the CHAMP1/POGZ/HP1α complex results in reduced H3K9me3 and loss of heterochromatin clustering. Interestingly, the POGZ subunit also binds and recruits the methyltransferase SETDB1 to sites of heterochromatin. Finally, peripheral blood lymphocytes from individuals with CHAMP1 syndrome also exhibit defective heterochromatin clustering and HR, suggesting that a defect in DNA repair underlies this syndrome. Taken together, the CHAMP1/POGZ/HP1α is a novel molecular machine which regulates heterochromatin clustering and contributes to local HDR repair in heterochromatin.

## RESULTS

### Structural modelling of the heterochromatic CHAMP1-POGZ-HP1α complex

Quantitative proteomic analyses of histone epigenetic marks and readers identified CHAMP1-POGZ-HP1α as a heterochromatin complex that binds H3K9me3^17^. Recent studies have shown that the CHAMP1^34,36^ and POGZ subunits^35^ promote HR-mediated DSB repair through interaction with REV7. Nevertheless, the relationship between the heterochromatin localization of the complex and HR is unclear.

To investigate the organization of the multi-subunit CHAMP1 complex, we performed *in silico* structural modeling of binary interactions among the subunits with AlphaFold2-Multimer (AF2)^42,43^. This allowed us to predict a high-confidence interaction between the CHAMP1 N-terminal ZnF domain (1-87 aa) and the structured C-terminus of POGZ (1021-1410 aa) (**Fig. 1a,b**). The interaction between CHAMP1 and POGZ is formed primarily by hydrophobic residues at the interface (**Fig. 1a**). Conversely, our modelling showed that the CHAMP1 C-terminal ZNF domain forms an extended β-sheet-type interaction with residues (16-22 aa) from the N-terminus of HP1α (**Extended Data Fig. 1a**). Furthermore, HP1α homodimerizes through its C-terminal chromodomains (**Extended Data Fig. 1b**), while the HPZ domain of POGZ interacts with the same C-terminal chromodomain of HP1α from the opposite side of the dimerization interface (**Extended Data Fig. 1c**). Since CHAMP1 binds to the same site on the N-terminus of HP1α as H3K9me3, HP1α homodimerization through its C-terminal chromodomain may allow a HP1α dimer to bind CHAMP1 and H3K9me3 simultaneously.

**Fig. 1.**
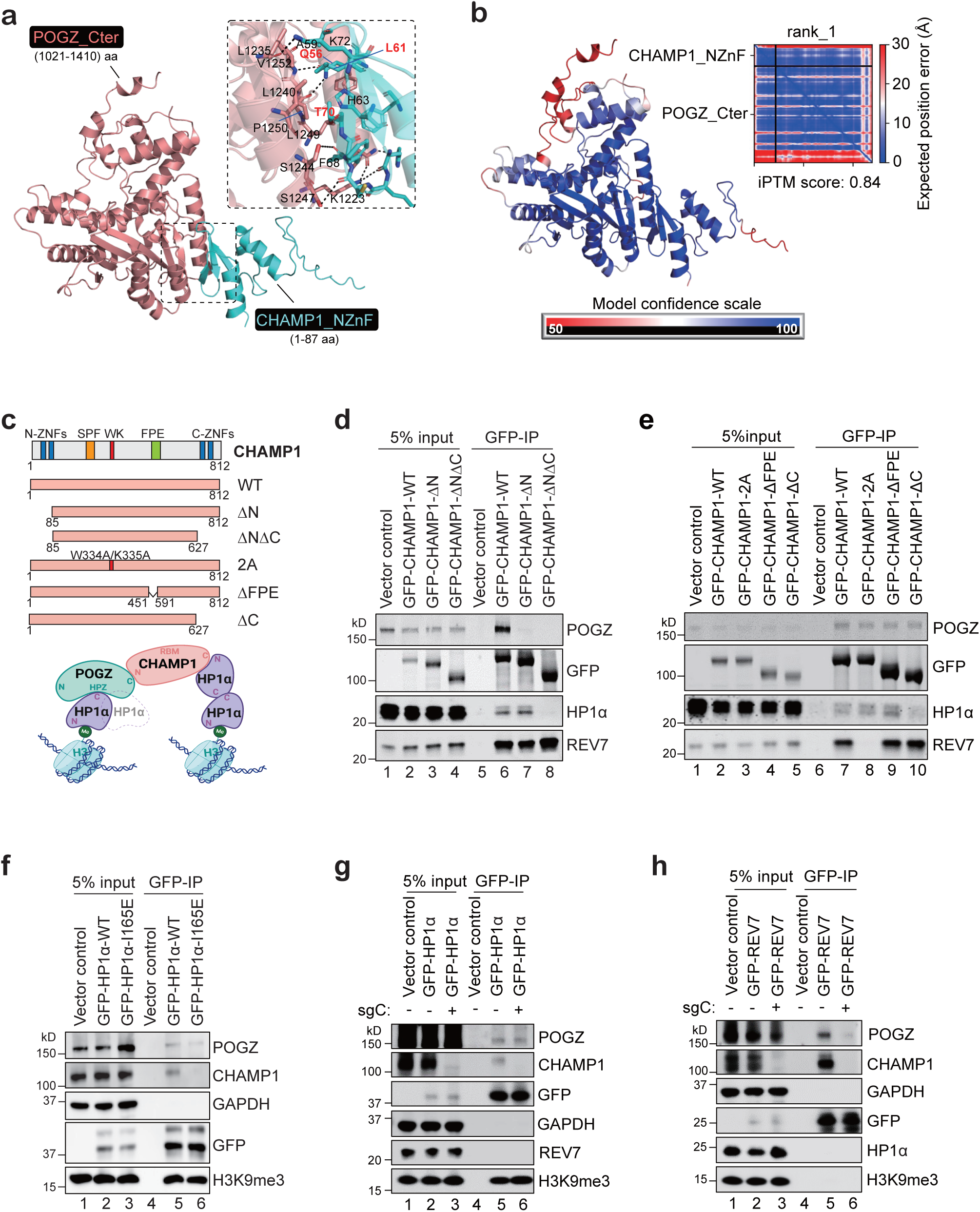
CHAMP1-POGZ-HP1α form a REV7 independent heterochromatin complex. **a.** AlphaFold2-Multimer (AF2)-predicted structural model of POGZ_Cter (peach)-CHAMP1_NZnF (cyan) complex. The key residues (shown in sticks) contributing to the protein-protein interactions are highlighted in the inset including CHAMP1 residues (Q56, L61 and T70) that are mutated to disrupt the interaction between POGZ and CHAMP1. **b**. The AF2-predicted model of POGZ_Cter-CHAMP1_NZnF is colored based on the confidence in the model prediction (100-high (blue) and 50-low (red)) and corresponding predicted aligned error plot (PAE) matrix showing the confidence in the predicted interaction between CHAMP1_NZnF and POGZ_Cter domains along with interface-predicted template scores (iPTM) for the AF2 prediction. **c.** (Top) Schematic of CHAMP1 wild-type (WT) and mutants. N-ZNFs (N-terminus C2H2-Zn finger domains), SPE (PxxSPExxK motifs), WK (SPxxWKxxP motifs), FPE (FPExxK motifs), C-ZNFs (C-terminus C2H2-Zn finger domains), and SBM (REV7 seatbelt-binding motif). 2A indicates W334A/K335A mutant. (Bottom) Schematic of CHAMP1 complex connecting two H3K9me3 sites. The N-terminus of CHAMP1 binds to the C-terminus of POGZ. Additionally, both the C-terminus of CHAMP and the HPZ motif of POGZ can independently bind to HP1α. CHAMP1 shows a preference for binding to the HP1α dimer, while POGZ is capable of binding to either the HP1α dimer or monomer. N-terminus of HP1α directly interacts with H3K9me3. **d**. Western blot showing GFP-immunoprecipitation of GFP-empty vector, GFP-CHAMP1 WT or N- or C-terminal deletion mutants, and the co-immunoprecipitation of endogenous POGZ, HP1α and REV7. **e.** Western blot showing GFP-immunoprecipitation of GFP-empty vector, GFP-CHAMP1 WT and GFP-CHAMP1 mutants in 293T cells, and the co-immunoprecipitation of endogenous POGZ, HP1α and REV7. **f**. Western blot showing GFP-immunoprecipitation of GFP-empty vector, GFP-HP1α WT or I165E mutant, and the co-immunoprecipitation of endogenous POGZ, CHAMP1 and H3K9me3. GAPDH acts as a negative control. **g.** Western blot showing GFP-immunoprecipitation of GFP-empty vector, GFP-HP1α in 293T WT and sgCHAMP1 cells, and the co-immunoprecipitation of endogenous POGZ, CHAMP1, REV7 and H3K9me3. GAPDH acts as negative control for IP. **h.** Western blot showing GFP-immunoprecipitation

Through these AF2 predictions and previously published studies on the interactions between POGZ and HP1α^19^, HP1α and H3K9me3 ^5^, and HP1α dimerization ^44,45^, we built a composite structural model of the CHAMP1 complex containing CHAMP1, POGZ, HP1α and H3K9me3 (**Fig. 1c**). In this model, the N-terminus of CHAMP1 interacts with POGZ, while the C-terminus of CHAMP1 directly engages with an HP1α dimer^5^. The CHAMP1-POGZ-HP1α complex appears to function as a bridge, connecting two H3K9me3. We next validated these predictions using a series of mutations in CHAMP1 and HP1α (homodimerization mutant) (**Fig. 1c-f, and Extended Data** Fig. 1d). As expected, N-terminal mutants of CHAMP1 (ΔN, ΔNΔC and QLT/RRR) failed to bind to POGZ (**Fig. 1d, lanes 7,8, and Extended Data** Fig. 1d**, lane 6**), and C-terminal mutants of CHAMP1 (ΔNΔC and ΔC) failed to bind to HP1α (**Fig. 1d, lane 8, and Fig. 1e, lane 10**). Furthermore, the HP1α dimerization mutant I165E^19,44^ failed to bind to CHAMP1, while POGZ could independently bind to HP1α and H3K9me3 (**Fig. 1f, lane 6**).

Interestingly, the GFP-CHAMP1-2A mutant, which failed to bind to REV7, still pulled down POGZ and HP1α (**Fig. 1e, lane 8**), suggesting that REV7 is not required for the formation of the CHAMP1/POGZ/HP1α heterochromatin complex. GFP-HP1α successfully pulled down CHAMP1, POGZ, and H3K9me3, but did not pull down REV7 (**Fig. 1g, lanes 5,6**). Moreover, GFP-REV7 pulled down CHAMP1 and POGZ but did not pull down HP1α or H3K9me3 (**Fig. 1h, lanes 5,6**). Taken together, our data suggest that the CHAMP1 complex may function in a REV7-independent pathway while interacting with heterochromatin.

### CHAMP1 complex promotes heterochromatin clustering at multiple genomic regions

To investigate the potential function of the CHAMP1 complex in the formation of heterochromatin clusters, we initially assessed the localization of CHAMP1 and POGZ within heterochromatin foci by immunofluorescence. Remarkably, we observed a strong colocalization of CHAMP1 and POGZ with H3K9me3, which we used as a marker of heterochromatin, in human U2OS and mouse NIH-3T3 cells. Colocalization was further enhanced by ionizing radiation (IR) induced DSBs (**Fig. 2a,b and Extended Data** Fig. 2a,b). Consistently, the association between CHAMP1 and HP1α was stronger after cellular exposure to IR (**Extended Data Fig. 2c**). This interaction was not affected by the loss of REV7 interaction (i.e., the CHAMP1-2A mutant), supporting the idea that the CHAMP1 heterochromatin complex functions independently of REV7. Interestingly, radiation also induced the interaction of H3K9me3 with HP1α, but not with HP1β or HP1γ (**Extended Data Fig. 2d**), suggesting a specific role of HP1α in the heterochromatin enrichment at DSBs.

**Fig. 2.**
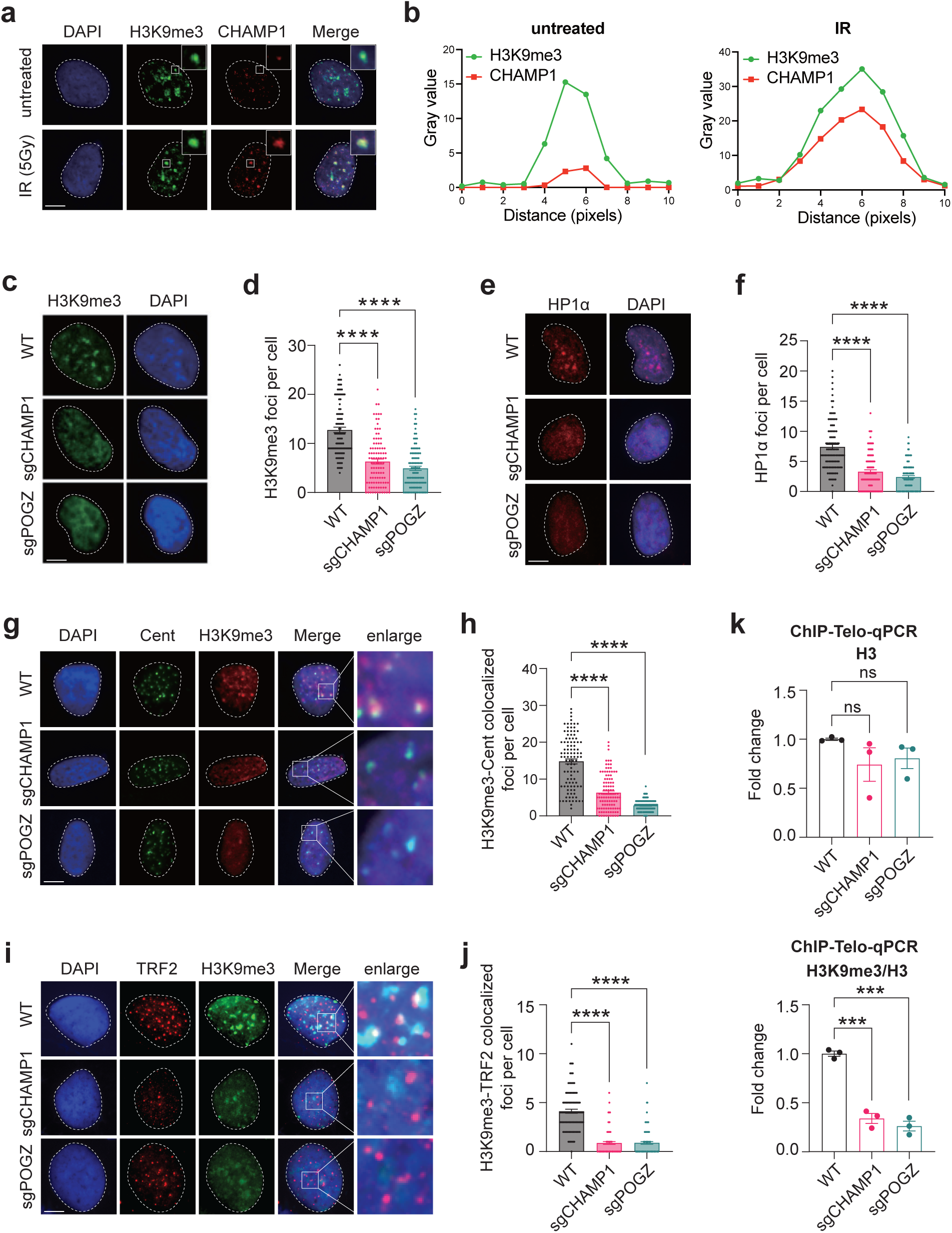
The CHAMP1 complex is required for the nuclear distribution of H3K9me3. **a.** gRepresentative immunofluorescence (IF) confocal images of the colocalized CHAMP1 and H3K9me3 foci with/without IR (5Gy) treatment in U2OS cells. DAPI was used to stain the Guclei. Scale bar, 5 µm. **b**. Square analysis of the intensity and colocalization of H3K9me3 and FCHAMP1 foci. **c**. Representative IF images of H3K9me3 foci in WT, sgCHAMP1 and sgPOGZ PU2OS cells. DAPI was used to stain the nuclei. Scale bar, 5 µm. **d**. Quantification of H3K9me3 foci in (c). Error bars indicate SEM. More than 100 cells were counted. ****P<0.0001. Statistical analysis was performed using two-tailed Student’s t test. **e**. Representative IF images omf HP1α foci in WT, sgCHAMP1 and sgPOGZ U2OS cells. DAPI was used to stain the nuclei. Scale bar, 5 µm. **f**. Quantification of HP1α foci in (e). Error bars indicate SEM. More than 100 cells were counted. ****P<0.0001. Statistical analysis was performed using two-tailed Student’s tntest. **g**. Representative IF-FISH confocal images of H3K9me3 colocalization with centromere (Cent) in WT, sgCHAMP1 and sgPOGZ U2OS cells. DAPI was used to stain the nuclei. Scale bar, 5 µm. **h**. Quantification of H3K9me3-Centromere colocalizations in (g). Error bars indicate SEM. More than 100 cells were counted. ****P<0.0001. Statistical analysis was performed using two-tailed Student’s t test. **i**. Representative IF confocal images of H3K9me3-TRF2 colocalization in WT, sgCHAMP1 and sgPOGZ U2OS cells. TRF2 was used as a telomere maker. DAPI was used to stain the nuclei. Scale bar, 5 µm. **j**. Quantification of H3K9me3 colocalization with TRF2 in (i). Error bars indicate SEM. More than 100 cells were counted. ****P<0.0001. Statistical analysis was performed using two-tailed Student’s t test. **k**. ChIP-qPCR analyses showing relative enrichment levels of H3 (Top) and H3K9me3 (Bottom) at telomeres. Level of H3K9me3 was normalized to total H3 levels (H3K9me3/H3). Error bars indicate SD. P-values calculated using Student’s T-test (***P<0.001; ns, non-significant).

Notably, the CRISPR-knockout (KO) of CHAMP1 or POGZ in U2OS cells resulted in a significant reduction in the formation of H3K9me3 and HP1α foci (**Fig. 2c-f**). However, the knockout of CHAMP1 or POGZ did not affect the overall expression level or chromatin binding of H3K9me3 and HP1α (**Extended Data Fig. 2e**,f**, lane 9-12**). To further determine whether the CHAMP1 heterochromatin complex promotes H3K9me3 accumulation at centromeres and telomeres, well-known heterochromatin sites, we next performed immunofluorescence with a centromere marker (Cent FISH) or a telomere marker (anti-TRF2 antibody)^46^. As expected, we observed a significant loss of H3K9me3 foci at both centromeres and telomeres following deletion of the CHAMP1 or POGZ in U2OS cells (**Fig. 2g-j).** Consistently, ChIP-qPCR also demonstrated a decreased level of H3K9me3 at telomeres in CHAMP1 or POGZ KO U2OS cells (**Fig. 2k**). Taken together, the CHAMP1 complex plays a role in assembling the H3K9me3 heterochromatin mark at multiple sites in the genome, including sites with damage-induced DSBs as well as at centromeres and telomeres.

Recent studies on telomeres in ALT tumor cells suggest a distinctive heterochromatin structure characterized by high levels of H3K9me3, low levels of ATRX, and elevated HDR processes at telomeres^47–49^. We therefore decided to investigate the possible role of the CHAMP1 complex in the maintenance of these ALT telomere structures and HDR repair. We first examined the possible localization of the CHAMP1 complex at telomeres in ALT U2OS cells.

Notably, we observed a strong colocalization of CHAMP1 and POGZ with telomere foci, particularly with large telomere foci, in U2OS cells (**Extended Data Fig.3a,b**). This suggests that the CHAMP1 complex may play a role in promoting the formation of telomere clusters. DSBs at telomeres are known to initiate long-range movements and clustering of telomeres, and these processes are critical for homology-directed telomere synthesis^50^. Accordingly, we evaluated the response of U2OS cells to engineered DSBs at telomeres, using the TRF1-FokI^50,51^. TRF1-FokI induced telomeric DSBs (**Extended Data Fig.3c,d**), resulting in telomere clustering (**Fig. 3a,b**), as measured by an increase in the average size of telomere foci in ALT cells and a decrease in the total number ^50^ (**Fig. 3c,d**). Interestingly, this DSB-induced telomere clustering was significantly diminished in CHAMP1 or POGZ knockout U2OS cells (**Fig. 3c,d**). Depletion of CHAMP1 or POGZ also led to a decrease in the hallmarks of ALT recombination, including a reduction in ALT-associated PML bodies (APBs)^14,52^ (**Fig. 3e,f**), which are sites of telomeric clustering and recombination^15,53^, and a decrease of non-S phase telomeric DNA synthesis^16^ (**Fig. 3g-i**). Taken together, our results demonstrate that the CHAMP1 complex plays a critical role in facilitating damage-induced telomere clustering and in maintaining ALT activity (**Fig. 3j**).

**Fig. 3.**
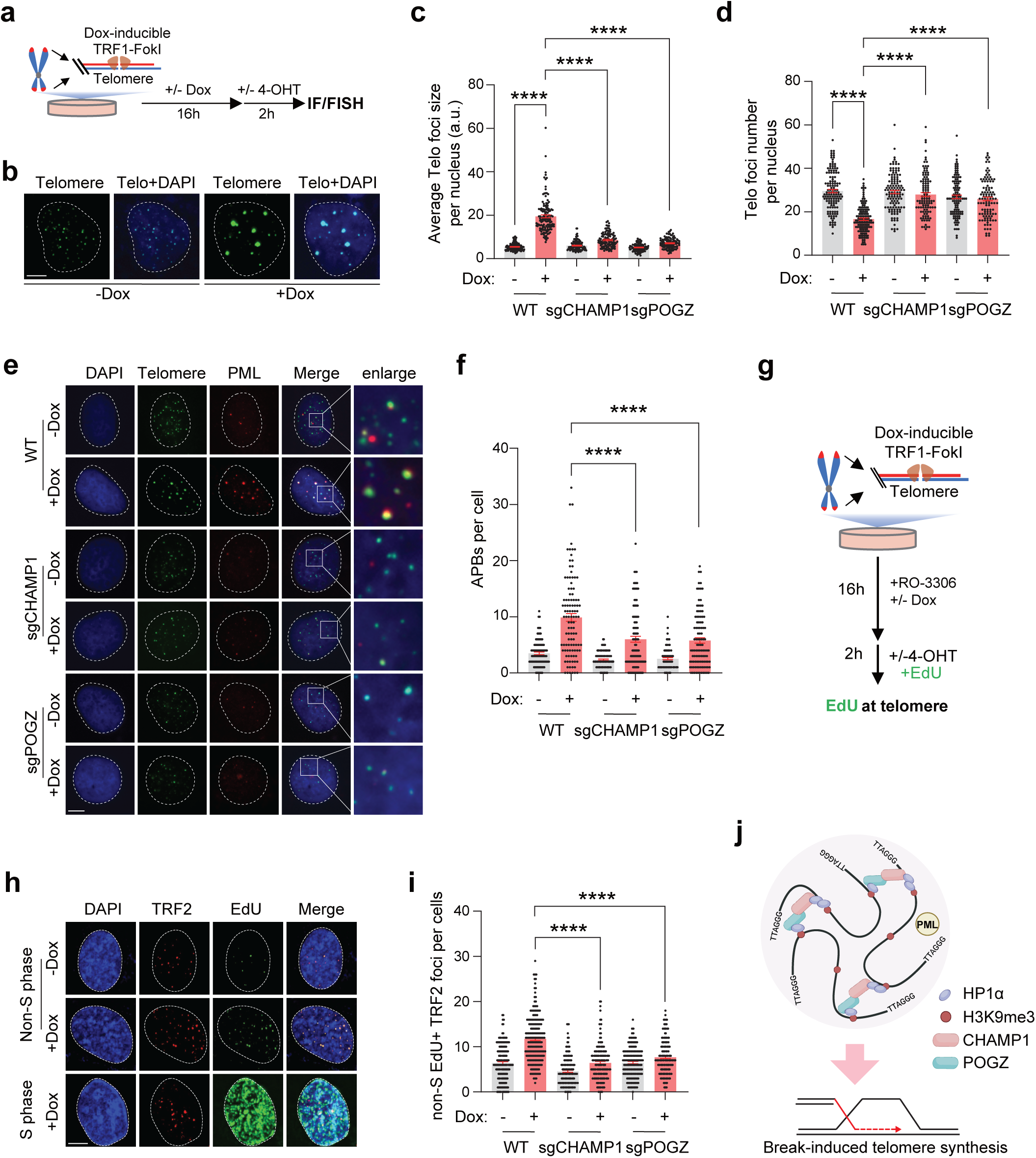
The CHAMP1 Complex promotes break-induced telomere clustering and synthesis. **a.** Schematic of U2OS-TRF1-FokI cell and time course of IF/FISH following Dox/4-OHT treatment. **b**. Representative FISH confocal images with telomere probe (green) treated with/without TRF1-FokI in U2OS cells. DAPI was used to stain the nuclei. Scale bar, 5 µm. **c-d**. CHAMP1 and POGZ KO U2OS cells were examined by telomere FISH after TRF1-FokI treatment (a). Average telomere (Telo) foci size (c) and telomere number (d) per nucleus was calculated using ImageJ. Error bars indicate SEM. More than 100 cells were counted. ****P<0.0001. Statistical analysis was performed using two-tailed Student’s t test. **e**. PML immunostaining combined with telomere FISH in wild-type (WT), sgCHAMP1 and sgPOGZ U2OS cells after TRF1-FokI treatment (a). Colocalization of PML foci and telomere signals (APBs) are shown as indicated. DAPI was used to stain the nuclei. Scale bar, 5 µm. **f**. Quantification of APBs shown in (e). Error bars indicate SEM. More than 100 cells were counted. ****P<0.0001. Statistical analysis was performed using two-tailed Student’s t test. **g**. Schematic of U2OS-TRF1-FokI cell and time course of IF assay following Dox/4-OHT and EdU treatment. **h**. Representative confocal images of immunostaining of TRF2 and EdU foci in S phase and non-S phase of U2OS cells following the treatment in (g). DAPI was used to stain the nuclei. Scale bar, 5 µm. **i**. Quantification of the colocalized TRF2 and EdU foci in WT, sgCHAMP1 and sgPOGZ U2OS cells. Error bars indicate SEM. More than 100 cells were counted. ****P<0.0001. Statistical analysis was performed using two-tailed Student’s t test. **j.** A cartoon showing that CHAMP1 complex promotes telomere clustering, APBs formation and facilitates Break-induced new telomeric DNA synthesis.

### CHAMP1 complex promotes HDR at ALT telomeres

We next used the PICh (<u>P</u>roteomics of <u>I</u>solated <u>Ch</u>romatin Segments) method to identify the recruitment of telomere-associated proteins following DSBs, in the presence or absence of the CHAMP1 heterochromatin complex (**Fig. 4a and Extended Data** Fig. 4a). Consistent with previous observations^51^, multiple DNA repair- and telomere maintenance-related biological processes were significantly upregulated at telomeres following TRF1-FokI induction (**Extended Data Fig. 4b**). CRISPR knockout of CHAMP1 resulted in a decrease of other CHAMP1 complex proteins (i.e., POGZ and HP1α) at telomeres with DSBs, as well as a decrease of many proteins associated with HR (e.g., RAD51, BRCA1, and BARD1), as measured by mass spectrometry (**Fig. 4b**). These results are consistent with the known role of CHAMP1 and POGZ in HR repair^34–36^. Additionally, CHAMP1 depletion also resulted in a reduction in proteins involved in break-induced replication (BIR) or break-induced telomere synthesis (BITS) ^51,54^ (i.e., POLD3, POLD4, RFC and PCNA) (**Fig. 4b**). Western blot analysis of PICh-purified telomeres revealed an enrichment of the CHAMP1 complex (i.e., CHAMP1 and HP1α) and BITS proteins (i.e., POLD3, RAD18 and PCNA) at telomeres in U2OS cells following TRF1-FokI induction, but a reduction in CHAMP1-depleted U2OS cells (**Fig. 4c, lanes 9-12**).

**Fig. 4.**
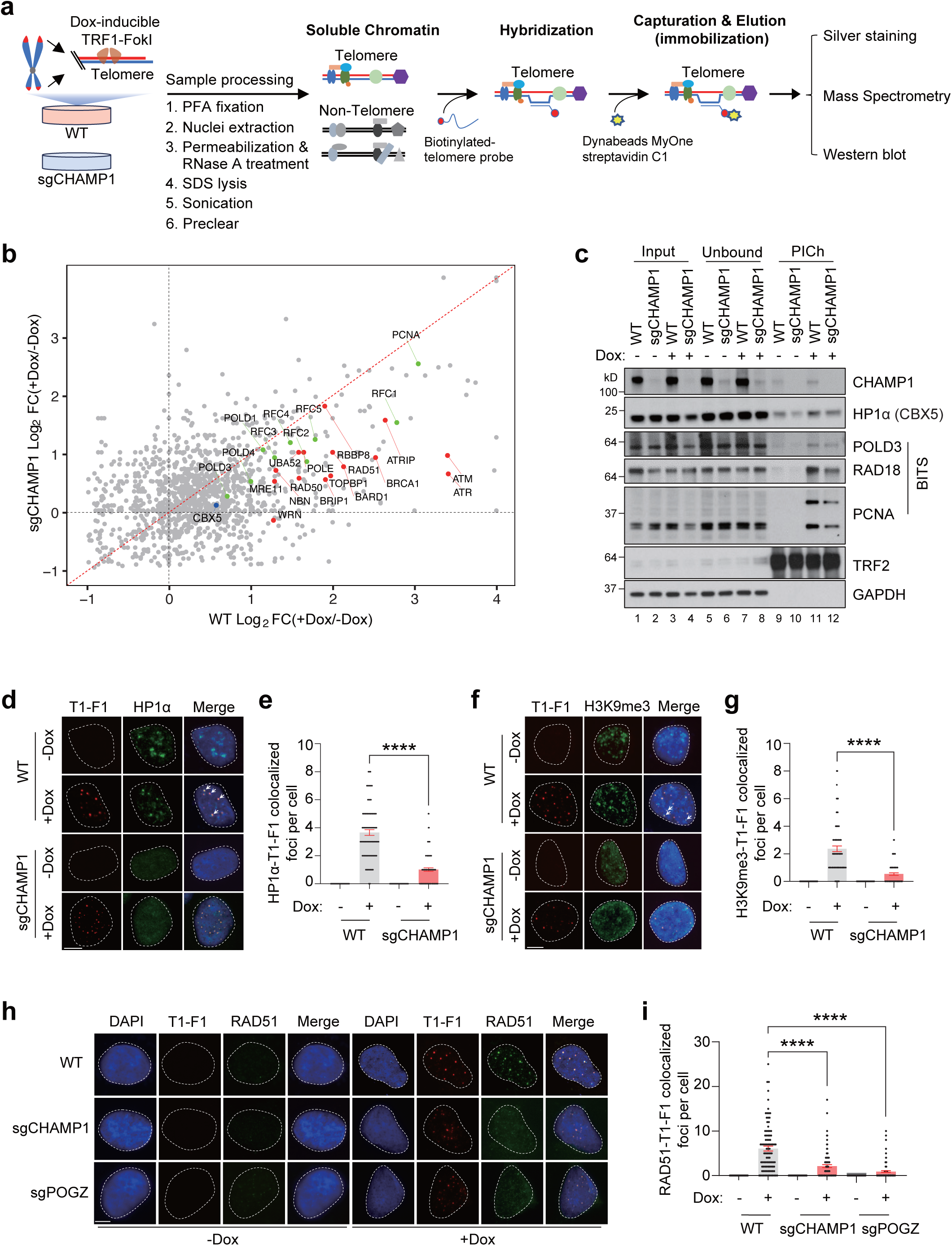
The CHAMP1 complex establishes telomeric heterochromatin and promotes the recruitment of HR and BIR factors. **a**. Schematic of the PICh approach used to define the telomere specific proteome in WT and sgCHAMP1 U2OS cells after TRF1-FokI (-/+Dox/4- OHT) treatment. Fractionated and precleared chromatin was hybridized to a biotinylated telomeric probe and subsequently captured on magnetic beads. Telomere-associated proteins were analyzed by silver staining, western blot analysis and mass spectrometry. PFA, paraformaldehyde. **b.** Scatterplot of the telomere specific DSB response proteome profile enriched by PICh in wild-type (WT) and sgCHAMP1 U2OS cells induced with TRF1-FokI. Mass Spectrometry data was shown in Table S1. Scatterplot of log2 FC[(+Dox+1)/(-Dox+1) of total peptide number] from two independent experiments. FC, fold change. Homologous recombination (HR) pathway proteins with Log2 FC > 1 in WT were highlighted in red. Break-induced replication (BIR) pathway proteins with Log2 FC > 1 in WT were highlighted in green. CBX5 (HP1α) was highlighted in blue. The red dotted line indicates the same value of log2 FC(+Dox/-Dox) in both WT and sgCHAMP1 cells. **c.** Western blot from PICh experiments in U2OS with/without Dox-induced TRF1-FokI. Telomere-associated proteins are blotted with the indicated antibodies. **d**. Representative IF confocal images depicting the colocalization of HP1α (Green) with TRF1-FokI (T1-F1, mCherry tagged, Red) in WT and sgCHAMP1 U2OS cells induced with/without TRF1-FokI for 2 hours. DAPI (blue) was used to stain the nuclei. Scale bar, 5 µm. **e**. Quantification of HP1α-TRF1-FokI colocalization events in (d). Error bars indicate SEM. More than 100 cells were counted. ****P<0.0001. Statistical analysis was performed using two-tailed Student’s t test. **f**. Representative IF confocal images depicting the colocalization of H3K9me3 (Green) with TRF1-FokI (T1-F1, mCherry tagged, Red) in WT and sgCHAMP1 U2OS cells induced with/without TRF1-FokI for 2 hours. DAPI (blue) was used to stain the nuclei. Scale bar, 5 µm. **g**. Quantification of H3K9me3-TRF1-FokI colocalization events in (f). Error bars indicate SEM. More than 100 cells were counted. ****P<0.0001. Statistical analysis was performed using two-tailed Student’s t test. **h**. Representative IF confocal images depicting the colocalization of RAD51 (Green) with TRF1-FokI (T1-F1, mCherry tagged, Red) in WT, sgCHAMP1 and sgPOGZ U2OS cells induced with/without TRF1-FokI for 2 hours. DAPI (blue) was used to stain the nuclei. Scale bar, 5 µm. **i**. Quantification of RAD51-TRF1-FokI colocalization events in (h). Error bars indicate SEM. More than 100 cells were counted. ****P<0.0001. Statistical analysis was performed using two-tailed Student’s t test.

Immunofluorescence further confirmed the reduction of CHAMP1 complex (i.e., POGZ and HP1α) and H3K9me3 at telomeres in CHAMP1 knockout U2OS cells (**Fig. 4d-g and Extended Data** Fig. 4c,d). Consistently, POGZ depletion also resulted in a reduction of proteins associated with HR and BITS at telomeres with DSBs (**Extended Data Fig. 4e**). Consistent with the local reduction in HR proteins, knockout of CHAMP1 or POGZ in U2OS cells resulted in a reduction in HR activity at telomeres, as measured by RAD51 and pRPA2(S33) foci at damaged telomeres (**Fig. 4h,i and Extended Data** Fig. 4f,g). Taken together, these results support a model in which the CHAMP1 complex promotes HDR, potentially through the clustering of heterochromatin at telomeres.

### CHAMP1 complex binds and recruits SETDB1 to telomeric heterochromatin clusters

To further evaluate the regulation of telomere heterochromatin deposition, we examined H3K9 methyltransferase levels and activity. Interestingly, SETDB1, but not other H3K9 methyltransferases, was recruited to telomeres after TRF1-FokI induction in U2OS cells, whereas this recruitment was not observed in CHAMP1- and POGZ-depleted U2OS cells (**Fig. 5a**). Immunofluorescence confirmed the reduction in SETDB1 at telomeres following TRF1-FokI induction in CHAMP1 or POGZ knockout U2OS cells (**Fig. 5b,c**). Based on previously published data, indicating a POGZ and SETDB1 interaction^55^, we reasoned that the CHAMP1 complex might directly recruit SETDB1 to telomeres. A strong interaction between SETDB1 and the Zn-fingers (type 2-7) of POGZ was predicted (iPTM =0.85) (**Fig. 5d**). Furthermore, an N-terminal deletion of CHAMP1, causing loss of the interaction with POGZ, also exhibited reduced interaction with SETDB1 (**Fig. 5e**). These results suggest that the CHAMP1 complex may recruit SETDB1 to telomeres through POGZ (**Extended Data Fig. 5a**).

**Fig. 5.**
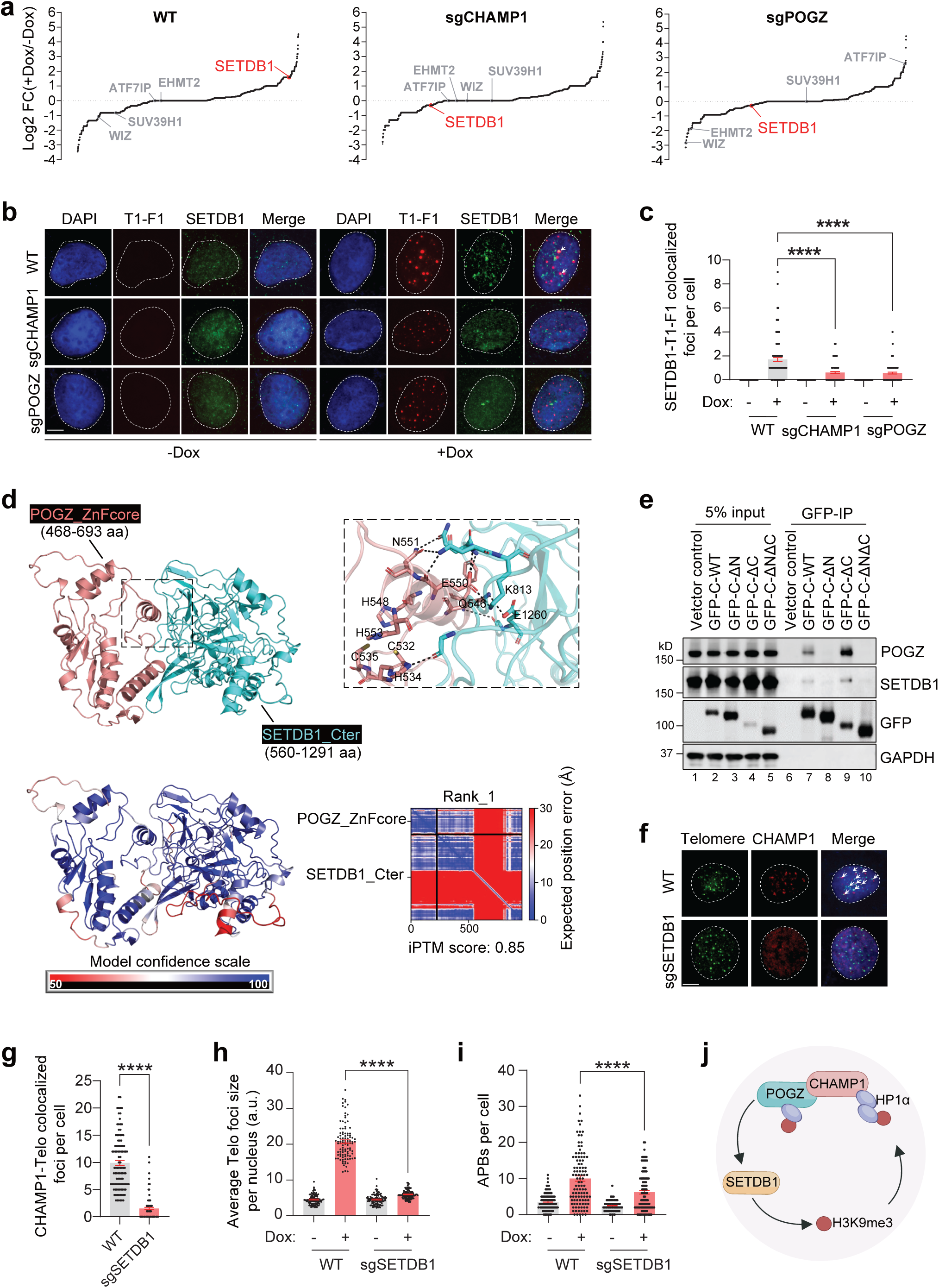
The CHAMP1 complex promotes the recruitment of SETDB1 to telomeres. **a.** Scatterplot of Log2 (+Dox+1)/(-Dox+1) of total peptide number from the telomere specific DSB response proteome profile enriched by PICh in wild-type (WT), sgCHAMP1 and sgPOGZ U2OS cells induced with TRF1-FokI. Mass Spectrometry data was shown in Table S2. SETDB1 is highlighted. **b**. Representative IF confocal images of SETDB1 (Green) colocalization with telomeres with double-strand breaks (T1-F1, mCherry, Red) in wild-type (WT), sgCHAMP1 and sgPOGZ U2OS cells induced with TRF1-FokI for 2 h. Arrow indicates SETDB1 and TRF1-FokI colocalization. DAPI was used to stain the nuclei. Scale bar, 5 µm. **c**. Quantification of (a) for number of SETDB1-mCherry colocalization events. Error bars indicate SEM. More than 100 cells were counted. ****P<0.0001. Statistical analysis was performed using two-tailed Student’s t test. **d.** (Top) AlphaFold2-Multimer (AF2)-predicted structural model of POGZ_ZnFcore (peach)-SETDB1_Cter (cyan) complex. The key residues (shown in sticks) contributing to the protein-protein interactions are highlighted in the inset including residues (C532, C535, H548 and H534) from POGZ_ZnFcore potentially forming a C2H2-type Zn finger that could be critical in maintaining interaction between POGZ and SETDB1. (Bottom) The AF2-predicted model of POGZ_ZnFcore-SETDB1_Cter complex is colored based on the confidence in the model prediction (100-high (blue) and 50-low (red)) and corresponding PAE matrix showing the confidence in the predicted interaction between POGZ_ZnFcore and SETDB1_Cter domains along with iPTM socres for the AF2 prediction. **e.** Western blot showing GFP-immunoprecipitation of GFP-vector control, GFP-CHAMP1 WT or deletion mutants, and the co-immunoprecipitation of endogenous SETDB1 and POGZ. GAPDH acts as a negative control for IP. **f.** Representative IF-FISH confocal images of CHAMP1 colocalization with telomeres in wild-type (WT) and sgSETDB1 U2OS cells. Arrow indicates CHAMP1-telomere colocalization. DAPI (blue) was used to stain the nuclei. Scale bar, 5 µm. **g**. Quantification of (f) for number of CHAMP1-telomere colocalization events. Error bars indicate SEM. More than 100 cells were counted. ****P<0.0001. Statistical analysis was performed using two-tailed Student’s t test. **h**. SETDB1 KO TRF1-FokI U2OS cells were examined by FISH using a FITC-labeled telomere probe after DOX/4-OHT treatment. Average telomere foci size per nucleus was calculated using ImageJ. Error bars indicate SEM. More than 100 cells were counted. ****P<0.0001. Statistical analysis was performed using two-tailed Student’s t test. **i**. PML immunostaining combined with telomere FISH in SETDB1 KO U2OS cells after DOX/4-OHT treatment. Colocalization of PML foci and telomere signals (APBs) are shown. Error bars indicate SEM. More than 100 cells were counted. ****P<0.0001. Statistical analysis was performed using two-tailed Student’s t test. **j.** A cartoon showing that the recruitment of SETDB1 by CHAMP1 complex could potentially facilitates a positive feedback loop that boosts the formation of telomere heterochromatin clusters.

Recently, it has been shown that ALT activity relies on a high level of H3K9 trimethylation, facilitated by SETDB1, and on the recruitment of recombination factors ^48^. Consistently, knockout of SETDB1 resulted in a decrease of H3K9me3 foci at telomeres (**Extended Data Fig. 5b**,c). Importantly, SETDB1 depletion decreased CHAMP1 recruitment to telomeres (**Fig. 5f,g**). Furthermore, SETDB1 depletion in U2OS cells led to reduced damage-induced telomere clustering and APBs (**Fig. 5h,i**). Collectively, these findings suggest that the CHAMP1 heterochromatin complex recruits SETDB1, thereby promoting a positive feedback loop that enhances the formation of telomere heterochromatin clusters (**Fig. 5j**).

### CHAMP1 and HP1α are epistatic in the regulation of heterochromatin clustering

Since HP1α is a critical subunit of the CHAMP1 heterochromatin complex, we next examined its role in the regulation of telomeric HR and clustering. Consistent with previous findings, HP1α is required for APBs formation^56^. HP1α knockdown resulted in a significant decrease of damage-induced telomere clustering and APBs formation (**Fig. 6a,b and Extended Data** Fig. 6a). HP1α knockdown did not result in a further decrease in damage-induced telomere clustering and APBs in CHAMP1 knockout U2OS cells (**Fig. 6a,b and Extended Data** Fig. 6a), indicating that CHAMP1 and HP1α are epistatic for APB formation and telomere clustering. Consistent with the requirement of HP1α dimerization in the function of the CHAMP1 complex, the wild-type HP1α protein, but not the HP1α-I165E dimerization mutant, restored telomere clustering and APBs in cells with an HP1α knockdown (**Fig. 6c,d and Extended Data** Fig. 6b). Furthermore, the C-terminal deletion mutant of CHAMP1, which lacks HP1α binding, and the N-terminus deletion mutant of CHAMP1, which lacks POGZ binding, did not restore RAD51 foci at telomeres or telomere clustering after CHAMP1 depletion (**Fig. 6e,f and Extended Data** Fig. 6c). These findings highlight the critical importance of the multi-subunit CHAMP1 heterochromatin complex in telomeric HR and clustering.

**Fig. 6.**
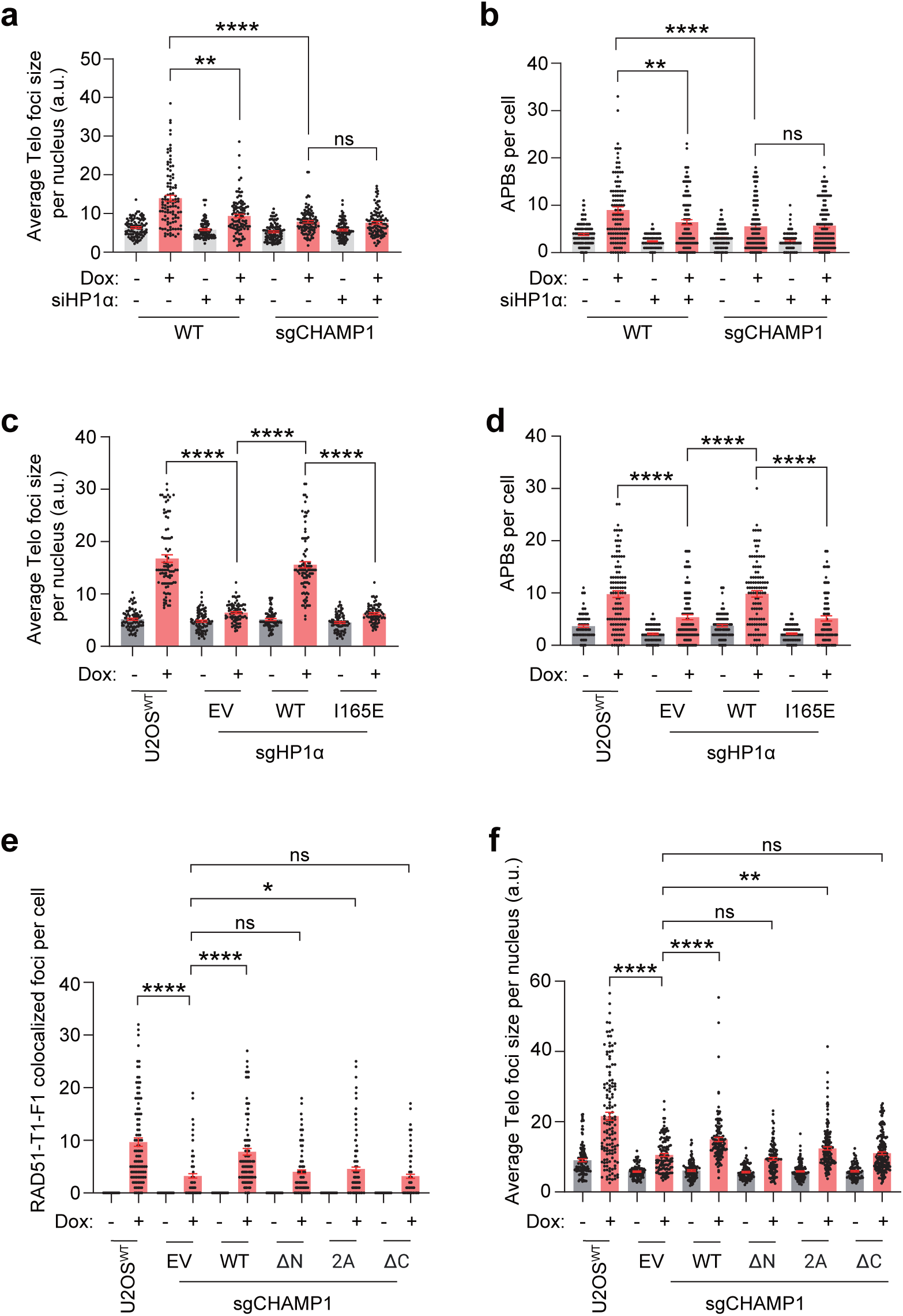
HP1α and CHAMP1 are epistatic in the regulation of telomeric HR and clustering. **a-b**. Wild-type (WT) and sgCHAMP1 U2OS-TRF1-FokI cells were treated with siControl or siHP1α, followed by examination of FISH using a telomere probe, or PML combined with telomere IF-FISH staining after DOX/4-OHT treatment. (a) Average telomere foci size per nucleus was calculated using ImageJ. Error bars indicate SEM. More than 100 cells were counted. ****P<0.0001, **P<0.01. Statistical analysis was performed using two-tailed Student’s t test. (b) Quantification of colocalization of PML foci and telomere signals (APBs). Error bars indicate SEM. More than 100 cells were counted. ****P<0.0001, **P<0.01. Statistical analysis was performed using two-tailed Student’s t test. **c-d.** WT, sgHP1α, and sgHP1α U2OS cells with ectopically expressed HP1α WT or mutant I165E were examined with FISH using a telomere probe, or PML combined with telomere IF-FISH staining using a FITC-labeled telomere probe after DOX/4-OHT treatment. (c) Average telomere foci size per nucleus was calculated using ImageJ. Error bars indicate SEM. More than 100 cells were counted. ****P<0.0001. Statistical analysis was performed using two-tailed Student’s t test. (d) Quantification of colocalization of PML foci and telomere signals (APBs). Error bars indicate SEM. More than 100 cells were counted. ****P<0.0001. Statistical analysis was performed using two-tailed Student’s t test. **e-f.** WT, sgCHAMP1, and sgCHAMP1 U2OS cells with ectopically expressed CHAMP1 WT or mutants (ΔN, 2A and ΔC) were examined with IF using anti-RAD51 or FISH using a telomere probe. (e) Quantification of RAD51-TRF1-FokI colocalization events. Error bars indicate SEM. More than 100 cells were counted. ****P<0.0001. Statistical analysis was performed using two-tailed Student’s t test. (f) Average telomere foci size per nucleus was calculated using ImageJ. Error bars indicate SEM. More than 100 cells were counted. ****P<0.0001, **P<0.01. Statistical analysis was performed using two-tailed Student’s t test.

### REV7 binding is not required for heterochromatin clustering or functional HR repair

The interaction of CHAMP1 and REV7 appears to be dispensable for the telomere HR and clustering (**Fig. 6e,f and Extended Data** Fig. 6c). REV7 does not co-immunoprecipitate with the CHAMP1 complex (**Fig. 1e,g,h**). Moreover, the CHAMP1-2A mutant protein, which has a disruption of the CHAMP1 binding site for REV7, can still assemble into the CHAMP1 complex, promote heterochromatin cluster formation, and promote HR activity (ie, RAD51 foci at sties of DSBs) (**Fig. 6e,f and Extended Data** Fig. 6c). Taken together, these results demonstrate that the activity of the REV7-independent CHAMP1 heterochromatin complex is essential for heterochromatic clustering and HR repair in heterochromatin.

### Patients with CHAMP1 syndrome display impaired heterochromatin clustering and defects in HR repair

We next examined EBV-immortalized lymphoblast cell lines derived from patients with CHAMP1 syndrome, resulting from an inherited mutation in one allele of the *CHAMP1* gene. The cells were obtained via the Coriell mutant cell line repository (**Fig. 7**). These cell lines express the mutant (truncated) CHAMP1 proteins (**Fig. 7a, and Extended Data** Fig. 7), as predicted by their genomic sequencing. These truncated proteins, lacking the C-terminal HP1α binding domain, may act as dominant negative proteins or lead to haploinsufficiency. Interestingly, these CHAMP1 patient-derived cell lines exhibit a defect in heterochromatin clustering, as indicated by a decrease of H3K9me3 foci (**Fig. 7b,c**) compared to wild-type control cells. The CHAMP1 patient-derived cell lines also have a defect in RAD51 foci assembly following IR (**Fig. 7d,e**), consistent with a deficiency in HR, and exhibit heightened sensitivity to both IR and PARP inhibitors (**Fig. 7f,g**). In summary, lymphocytes from individuals carrying CHAMP1 mutations exhibit a defect in heterochromatin clustering and deficiencies in homologous recombination, providing a possible functional diagnostic assay for the CHAMP1 syndrome.

**Fig. 7.**
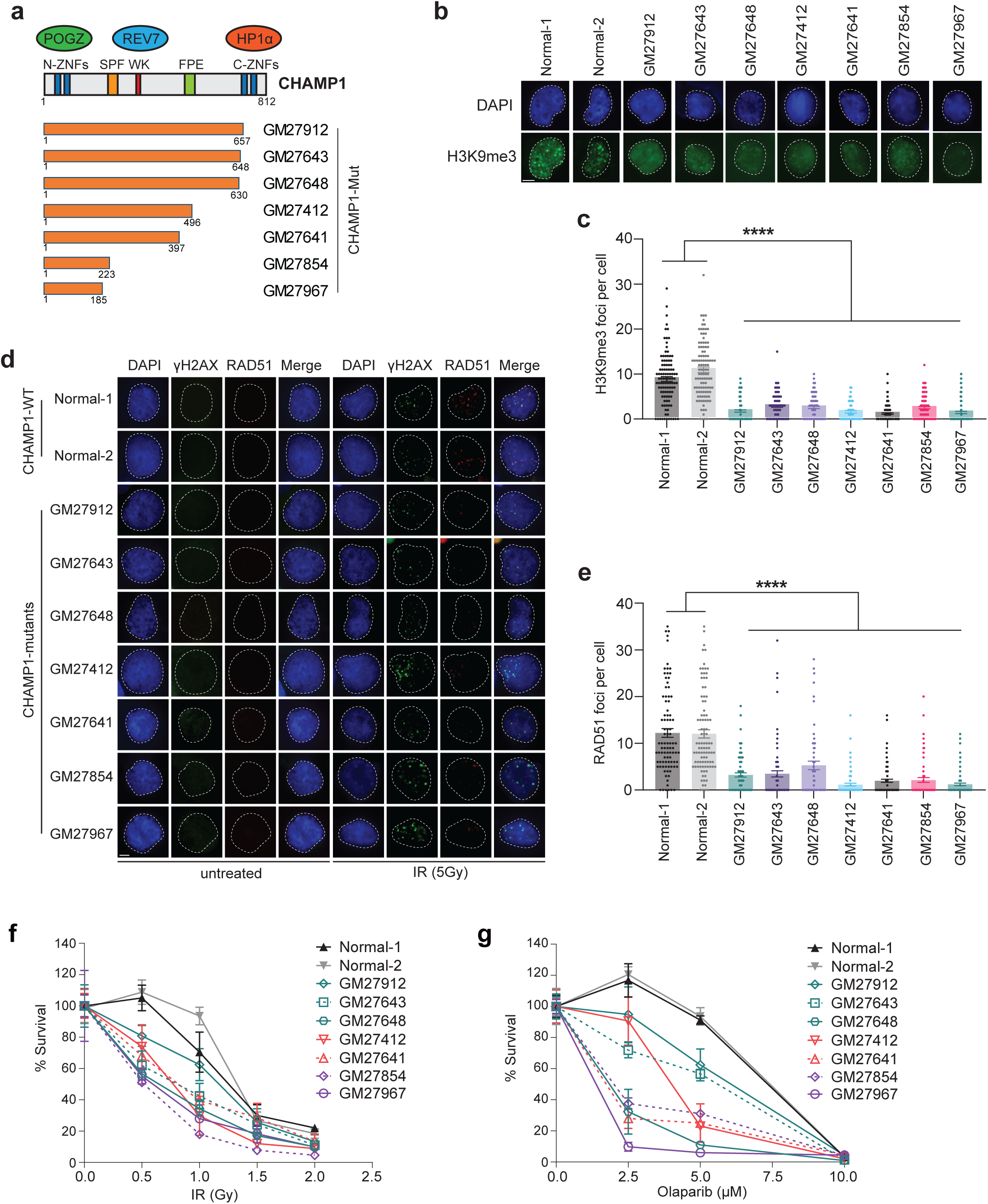
Peripheral blood lymphocytes from Patients with CHAMP1 syndrome display impaired heterochromatin clustering and defects in HR repair. **a.** Schematic of CHAMP1 mutants from patients with CHAMP1 syndrome. b. EBV-immortalized lymphoblast cell lines derived from several CHAMP1 patients (Coriell) were examined by immunostaining with H3K9me3 antibody. Representative images show the H3K9me3 foci. DAPI was used to stain the nuclei. Scale bar, 5 µm. c. Quantification of H3K9me3 foci of the CHAMP1 patient lymphoblast cells in (b). More than 100 cells were counted. ****P<0.0001. Statistical analysis was performed using one-way ANOVA. d. Representative images of RAD51 foci formation in normal and CHAMP1 patient lymphoblast cells following with/without 5Gy IR treatment. DAPI was used to stain the nuclei. Scale bar, 5 µm. e. Quantification of RAD51 foci of the CHAMP1 patient lymphoblast cells following with 5Gy IR treatment. More than 100 cells were counted. ****P<0.0001. Statistical analysis was performed using one-way ANOVA. f. 3-day cytotoxicity analysis of the normal and CHAMP1 patient lymphoblast cells treated with various dose of radiation. Cell viability was detected by CellTiter Glo (Promega). Error bars indicate SD, n=3 independent experiments. g. 3-day cytotoxicity analysis of the normal and CHAMP1 patient lymphoblast cells treated with various dose of Olaparib. Cell viability was detected by CellTiter Glo (Promega). Error bars indicate SD, n=3 independent experiments.

## DISCUSSION

Taken together, our studies show that the CHAMP1 heterochromatin complex, consisting of CHAMP1, POGZ, and HP1α, has a direct role in heterochromatin assembly and Homology-Directed DNA Repair. The complex binds to H3K9me3 in heterochromatin sites, including centromeres and telomeres, and contributes to the H3K9 trimethylation at these sites. The complex is also recruited to heterochromatic sites with DSBs generated by ionizing radiation. Moreover, the CHAMP1 heterochromatin complex plays a role in HDR at these heterochromatic sites.

Our results support the presence of two distinct CHAMP1/POGZ complexes. One is REV7-dependent, and the other is REV7-independent. We previously showed that the CHAMP1/POGZ complex promotes HR by binding to REV7 and limiting the binding of REV7 to proteins in the Shieldin complex (i.e., the REV7, SHLD1, SLD2, SHLD3 complex). The Shieldin complex is known to block the resection of DSB double strand breaks and thereby limit the level of HR repair. While this HR mechanism primarily occur in euchromatic regions, it may also occur to some extent in heterochromatic regions. However, in the current study, we identify another CHAMP1 complex, the CHAMP1 heterochromatin complex, whose function was previously unknown. This latter complex is enriched in heterochromatin and excludes the CHAMP1 binding partner REV7. Here, we show that this complex interacts directly with HP1α and promotes the H3K9me3 marks in heterochromatin. Moreover, this complex includes the SETDB1 methyltransferase. The complex is required for the recruitment of HDR proteins to heterochromatic regions of the chromosome. Loss of the CHAMP1 heterochromatin complex results in a decrease in repair of TRF1-FokI mediated DSBs at telomeres. Knockdown of CHAMP1 complex subunits, or mutation of individual subunits preventing complex assembly, reduced pRPA and RAD51 foci at these sites.

Evaluation of the CHAMP1-2A mutant protein, which fails to bind to REV7^36^ is especially informative. CHAMP1-2A protein does not enable HDR repair by preventing the binding of REV7 with the Shieldin complex^36^. However, the CHAMP1-2A mutant protein is still functional for HDR repair through its interaction with the CHAMP1 heterochromatin complex. As a subunit of the CHAMP1 heterochromatin complex, the CHAMP1-2A mutant protein still allows heterochromatin clustering, recruitment of HDR proteins to heterochromatin, and repair of DSBs in heterochromatin (i.e., formation of RAD51 foci).

Our study also demonstrates that the CHAMP1 heterochromatin complex is especially important to heterochromatin clustering and HDR repair at specific heterochromatic sites. For instance, the telomeres of ALT tumor cells are known to have high levels of H3K9me3, high HDR activity, and high levels of replication stress. ALT telomeres appear to have a unique heterochromatin environment ^48,57^. The CHAMP1 heterochromatin complex is required for assembly and maintenance of ALT telomeres, and knockdown or knockout of the CHAMP1 complex results in decreased DSB-induced telomere cluster formation and reduced HDR activity, coincident with a loss of telomere-associated histone H3 lysine 9 trimethylation (H3K9me3).

The CHAMP1 heterochromatin complex may play additional roles in heterochromatin structures. For instance, the complex may be regulated by the ATM kinase^2^ or may function in HDR repair at the periphery of the cell nuclease^9,18^. The complex may play a role in specific signaling events required for assembling heterochromatin at stressed replication forks^58,59^. Future studies will be required to assess the role of the CHAMP1 heterochromatin complex at these other heterochromatic sites.

A reduction in H3K9me3 is known to result from a mislocalization or inactivation of the SETDB1, a methyltransferase known to trimethylate H3K9^60^. Consistent with these findings, our AF2 analysis, immunoprecipitation studies, and PICh results revealed an interaction of SETDB1 with the POGZ subunit of the complex, required for the recruitment of SETDB1 to sites of heterochromatin. Taken together, the CHAMP1 complex appears to be a critical determinant of heterochromatin clustering. Genetic ablation of a subunit of the complex, or a mutation in a subunit which disrupts complex assembly, results in failure to recruit SETDB1 and in loss of heterochromatin clustering.

Previous studies have suggested that tightly-packed heterochromatin regions have an overall decrease in HDR activity^3^. Paradoxically, in our studies and in other recent reports^48,61,62^, the increase in H3K9 trimethylation or other modifications in the heterochromatin composition at telomeres, may enhance HDR activity. Consistent with these findings, the CHAMP1 complex may promote HDR by functioning as a bridge between two heterochromatin nucleosomes, either intra- or interchromosomally, through its interaction with H3K9me3 in heterochromatin. This bridge may facilitate the formation of heterochromatin clusters, thereby creating an environment that promotes the recruitment and stabilization of HDR proteins.

Finally, individuals with *de novo* heterozygous mutations in *CHAMP1*, known as CHAMP1 syndrome, present with a neurodevelopmental disorder characterized by intellectual disability, behavioral symptoms, and distinct dysmorphic features. Interestingly, *de novo* heterozygous mutations in the *POGZ* gene also result in another rare but highly-related neurodevelopment syndrome called the White Sutton Syndrome (WSS) ^25,27,28,30,31,63^. The mutant proteins encoded by pathogenic alleles of CHAMP1 have C-terminus truncations, and perhaps dominant negative activity, and cannot assemble into functional CHAMP1 complexes. Accordingly, lymphoblast cultures derived from individuals with CHAMP1 syndrome and expressing these mutant proteins have defects in heterochromatin and homologous recombination. These functional defects may also underlie the neurodevelopmental defects in these individuals. Lymphoblasts from these individuals have phenotypic characteristics similar to engineered human cells lacking the CHAMP1 complex. Taken together, these cellular characteristics could serve as a functional diagnostic screening tool for identifying individuals with CHAMP1 syndrome among the much larger number of individuals with autism spectrum disorders. In conclusion, we provide evidence that the CHAMP1 heterochromatin complex is critical for the heterochromatin HDR repair, not only in tumor cells but also in normal tissue cells. The clinical significance of the CHAMP1 complex will require further studies.

