## Extended Data Figures and Figure legends for "CHAMP1 complex directs heterochromatin assembly and promotes homology-directed DNA repair": Extended Data Figure Legends.docx

**Extended Data Fig. 1, related to Fig. 1. CHAMP1, POGZ, and HP1α combine to create a complex. a.** AF2-predicted structural model of CHAMP1_CZnF (peach)-HP1α_Nter (cyan). The key residues (shown in sticks) contributing to the formation of extended 𝛽-sheet interactions between CHAMP1 and HP1α are highlighted in the inset. The AF2-predicted model of CHAMP1_CZnF-HP1α_Nter complex is colored based on the confidence in the model prediction (100-high (blue) and 50-low (red)) and corresponding PAE matrix showing the confidence in the predicted interaction between CHAMP1_CZnF and HP1α_Nter along with iPTM score for the AF2 prediction. **b.** AlphaFold2-Multimer (AF2)-predicted structural model of homodimerization of HP1α_Cter. The key residues (shown in sticks) contributing to the protein-protein interactions are highlighted in the inset including HP1α residue (I165) that is shown to disrupt the homodimerization of HP1α when mutated. The AF2-predicted model of HP1α_Cter dimer is colored based on the confidence in the model prediction (100-high (blue) and 50-low (red)) and corresponding PAE matrix showing the confidence in the predicted interaction between two HP1α_Cter domains along with iPTM score for the AF2 prediction. **c.** AF2-predicted structural model of POGZ_HPZ (peach)-HP1α_Cter (cyan). The key residues (shown in sticks) contributing to the protein-protein interactions are highlighted in the inset including residues (C817, C820, H833 and H840) from POGZ_HPZ forming a C2H2-type Zn finger which was shown to be critical to maintain interactions with HP1α. The AF2-predicted model of POGZ_HPZ- HP1α_Cter complex is colored based on the confidence in the model prediction (100-high (blue) and 50-low (red)) and corresponding PAE matrix showing the confidence in the predicted interaction between POGZ_HPZ motif and HP1α_Cter domain along with iPTM score for the AF2 prediction. **d.** Western blot showing GFP-immunoprecipitation of GFP-empty vector, GFP-CHAMP1-wild-type (WT) and GFP-CHAMP1-QLT/RRR mutant in 293T cells, and the co-immunoprecipitation of endogenous POGZ and REV7. GAPDH acts as negative control for IP.

**Extended Data Fig. 2, related to Fig. 2. IR induces CHAMP1-POGZ-HP1α complex formation. a.** Representative immunofluorescence (IF) confocal images of the colocalized CHAMP1 and POGZ at DAPI dense heterochromatin clusters with/without IR (5Gy) treatment in mouse NIH-3T3 cells. DAPI was used to stain the nuclei. Scale bar, 5 µm. **b**. Square analysis of the intensity and colocalization of DAPI, CHAMP1 and POGZ foci in (a). **c.** (Left) Western blot showing GFP-immunoprecipitation of GFP-tagged CHAMP1-WT and -2A mutant in 293T cells after IR (5Gy) treatment, and the co-immunoprecipitation of endogenous REV7 and HP1α. (Right) The bands of HP1α after IP was calculated using ImageJ. Error bars indicate SEM. **P<0.01, ***P<0.001, ns, non-significant. Statistical analysis was performed using two-tailed Student’s t test. **d**. Western blot showing GFP-immunoprecipitation of GFP-tagged HP1β, HP1γ, HP1α in 293T cells after IR (5Gy) treatment, and the co-immunoprecipitation of endogenous H3K9me3 and γH2AX. **e**. Western blot showing the cytoplasmic (Cyto), soluble nuclear (S-Nuc) and chromatin fraction in wild-type (WT) and sgCHAMP1 U2OS and HeLa cells. **f.** Western blot showing the cytoplasmic (Cyto), soluble nuclear (S-Nuc) and chromatin fraction in WT and sgPOGZ U2OS and HeLa cells. All of the immunoblots are representative of at least two independent experiments.

**Extended Data Fig. 3, related to Fig. 3. CHAMP1 complex binds to the large ALT telomeres. a-b**. IF-FISH confocal analysis of the colocalization of CHAMP1 (a) or POGZ (b) and telomere using a telomere (Telo) probe (green). DAPI was used to stain the nuclei. Scale bar, 2 µm. Telomere size was calculated using ImageJ. Error bars indicate SEM. ****P<0.0001. Statistical analysis was performed using two-tailed Student’s t test. **c**. The mCherry-tagged TRF1-FokI colocalizes with the shelterin protein TRF2. Representative IF confocal images of telomeric binding protein TRF2 (green) and TRF1-FokI (T1-F1, mCherry, red) in the U2OS-TRF1-FokI cells with/without Dox/4-OHT treatment. DAPI was used to stain the nuclei. Scale bar, 5 µm. **d**. Representative IF confocal images of telomeric binding protein TRF2 (red) and γH2AX (green) in the U2OS-TRF1-FokI cells with/without Dox/4-OHT treatment. DAPI was used to stain the nuclei. Scale bar, 5 µm.

**Extended Data Fig. 4, related to Fig. 4. CHAMP1 complex promotes DSB HR and BIR repair at telomeres. a.** Silver staining of telomere chromatin bound fractions enriched by PICh with telomeric probe in wild-type (WT) and sgCHAMP1 U2OS cells induced with/without TRF1-FokI for 2 hrs. **b.** Pathway analysis of the telomere-specific DSB response proteome enriched by PICh in WT U2OS cells. Mass Spectrometry data was shown in Table S1. **c**. Representative IF confocal images showing the colocalization of POGZ (Green) with TRF1-FokI (mCherry, Red) in WT and sgCHAMP1 U2OS cells induced with TRF1-FokI for 2 h. DAPI was used to stain the nuclei. Scale bar, 5 µm. **d**. Quantification of (c) for number of POGZ-mCherry colocalization events. Error bars indicate SEM. More than 100 cells were counted. ****P<0.0001. Statistical analysis was performed using two-tailed Student’s t test. **e.** Scatterplot of the telomere specific DSB response proteome profile enriched by PICh in wild-type (WT) and sgPOGZ U2OS cells induced with TRF1-FokI. Mass Spectrometry data was shown in Table S2. Scatterplot shows Log2 FC[(+Dox+1)/(-Dox+1) of total peptide number]. FC, fold change. Homologous recombination (HR) pathway proteins with Log2 FC > 0.5 in WT were highlighted in red. Break-induced replication (BIR) pathway proteins with Log2 FC > 0.5 in WT were highlighted in green. The red dotted line indicates the same value of log2 FC(+Dox/-Dox) in both WT and sgCHAMP1 cells. **f**. Representative IF images depicting the colocalization of pRPA2(S33) (Green) with TRF1-FokI (mCherry, Red) in WT, sgCHAMP1 and sgPOGZ U2OS cells induced with/without TRF1-FokI for 2 hours. DAPI was used to stain the nuclei. Scale bar, 5 µm. **g**. Quantification of pRPA2(S33)-mCherry colocalization events in (f). Error bars indicate SEM. More than 100 cells were counted. ****P<0.0001. Statistical analysis was performed using two-tailed Student’s t test.

**Extended Data Fig. 5, related to Fig. 5. CHAMP1 complex recruits SETDB1 to promote heterochromatin formation at telomeres. a.** AlphaFold2-Multimer (AF2)-predicted structural model showing the interaction of SETDB1 and the CHAMP1 complex. **b.** Immunoblotting against SETDB1 in WT and sgSETDB1 U2OS cells. GAPDH was used as a loading control. **c**. Quantification of H3K9me3-TRF2 colocalization in WT and sgSETDB1 U2OS cells. Error bars indicate SEM. More than 100 cells were counted. ****P<0.0001. Statistical analysis was performed using two-tailed Student’s t test.

**Extended Data Fig. 6, related to Fig. 6. CHAMP1 complex subunit interaction is crucial for ALT activity. a.** Immunoblotting against CHAMP1 and HP1α in WT and sgCHAMP1 U2OS cells after siRNA treatment against HP1α. GAPDH was used as a loading control. **b**. Immunoblotting against HP1α was performed in WT, sgHP1α, and sgHP1α U2OS cells with ectopically expressed HP1α WT or mutant I165E. Vinculin was used as a loading control. **c**. Immunoblotting against GFP in WT, sgCHAMP1, and sgCHAMP1 U2OS cells with ectopically expressed CHAMP1 WT or mutants (ΔN, 2A and ΔC). GAPDH was used as a loading control.

**Extended Data Fig. 7, related to Fig. 7. Lymphocyte cells from CHMAP1 patients express truncated CHAMP1 proteins.** Western blot of CHAMP1 in EBV-immortalized lymphoblast cell lines derived from several CHAMP1 patients (Coriell). GAPDH acts as loading control.
