## Extended Data Figures and Figure legends for "CHAMP1 complex directs heterochromatin assembly and promotes homology-directed DNA repair": Extended Data Figures.pdf

**Extended Data Fig. 1**

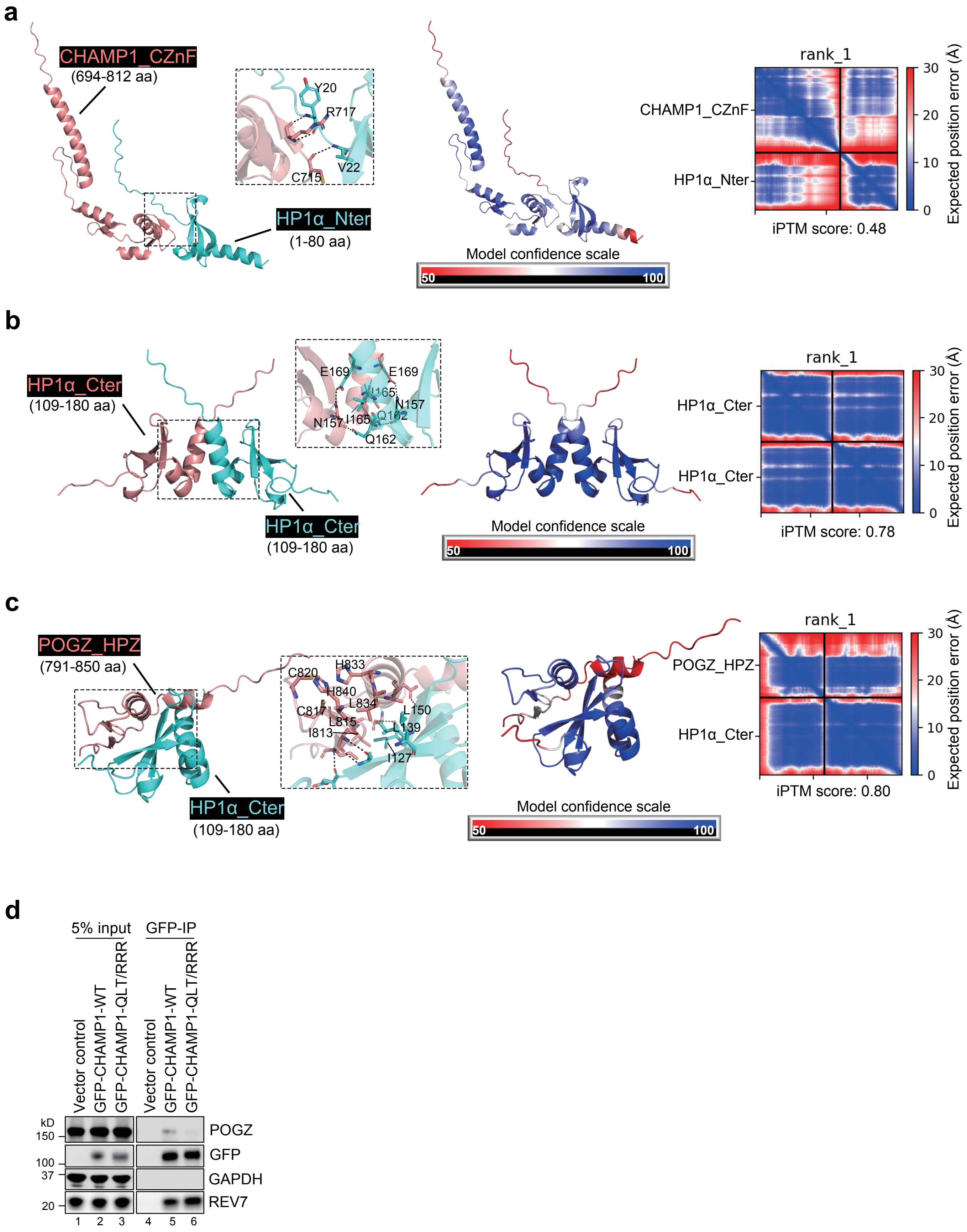

### Extended Data Fig. 2

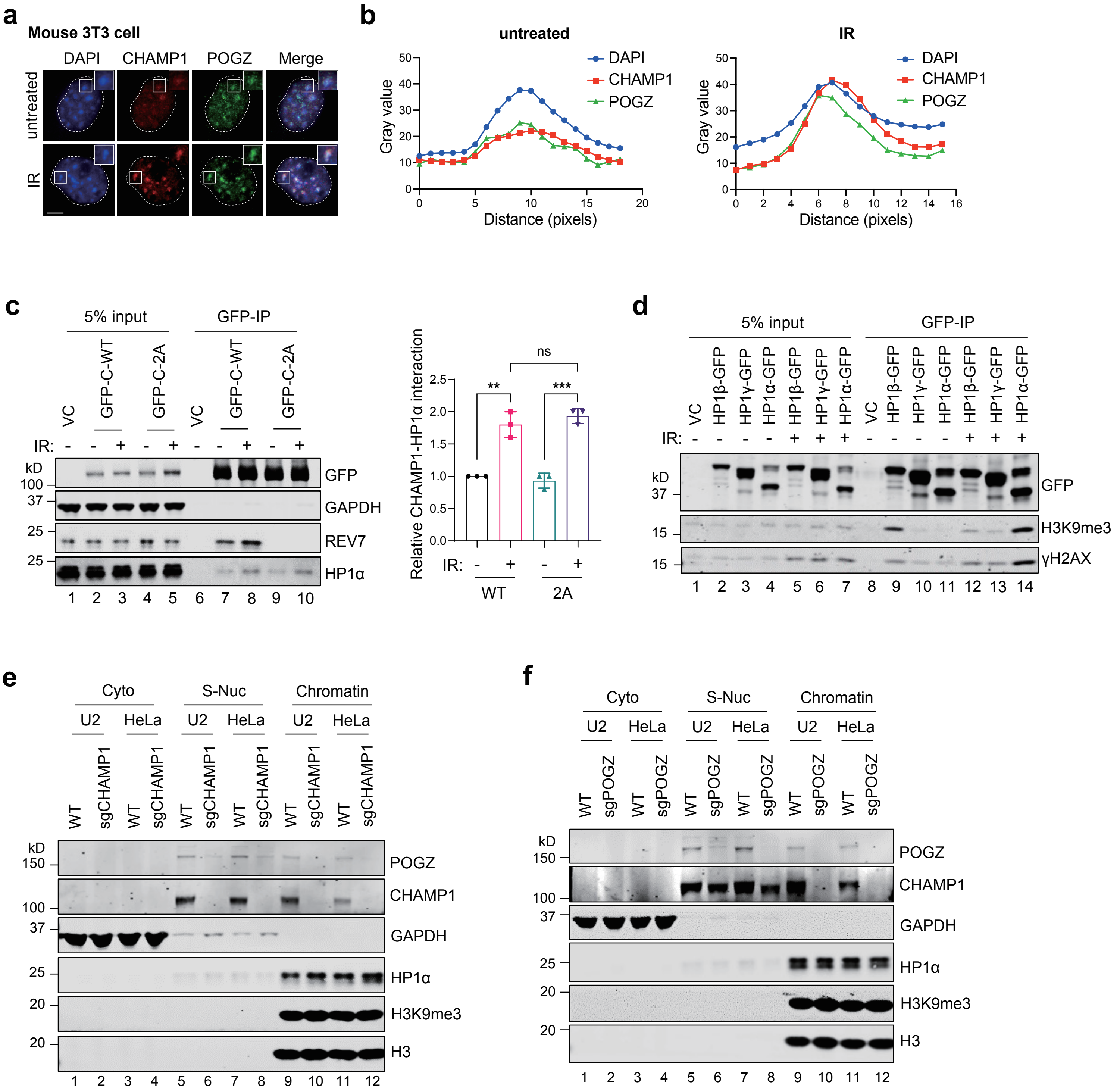

### Extended Data Fig. 3

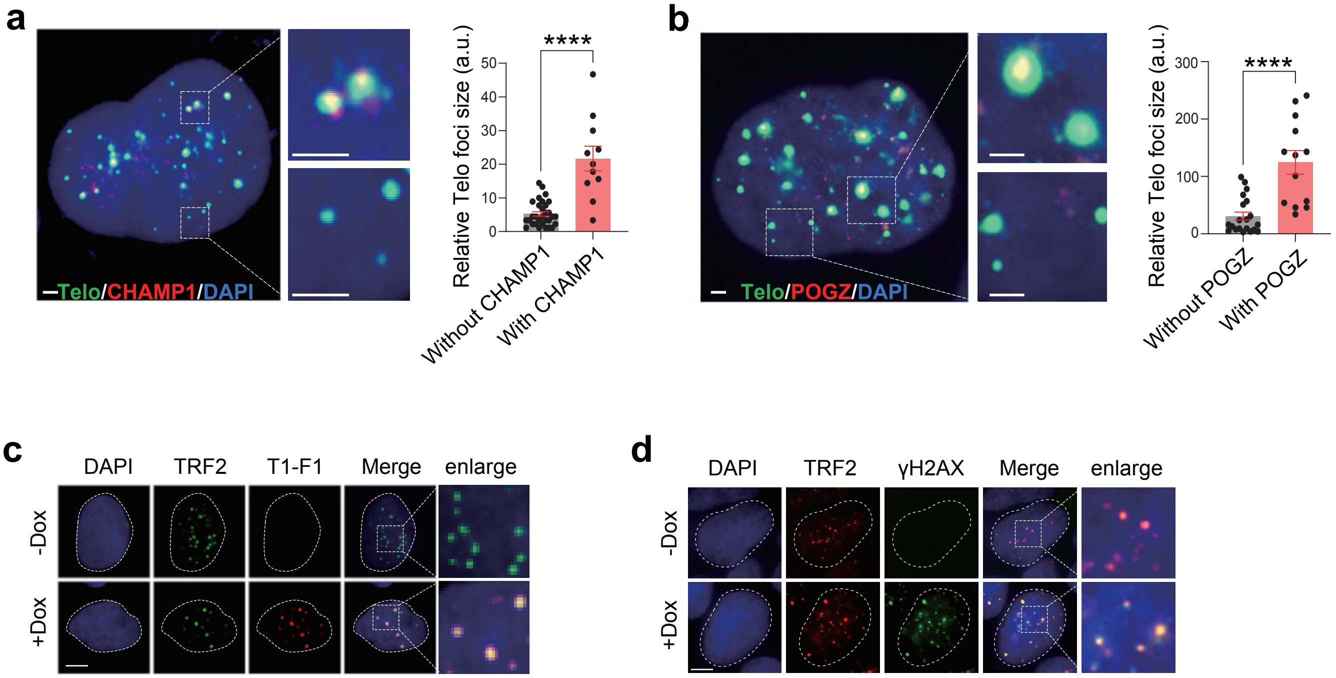

### Extended Data Fig. 4

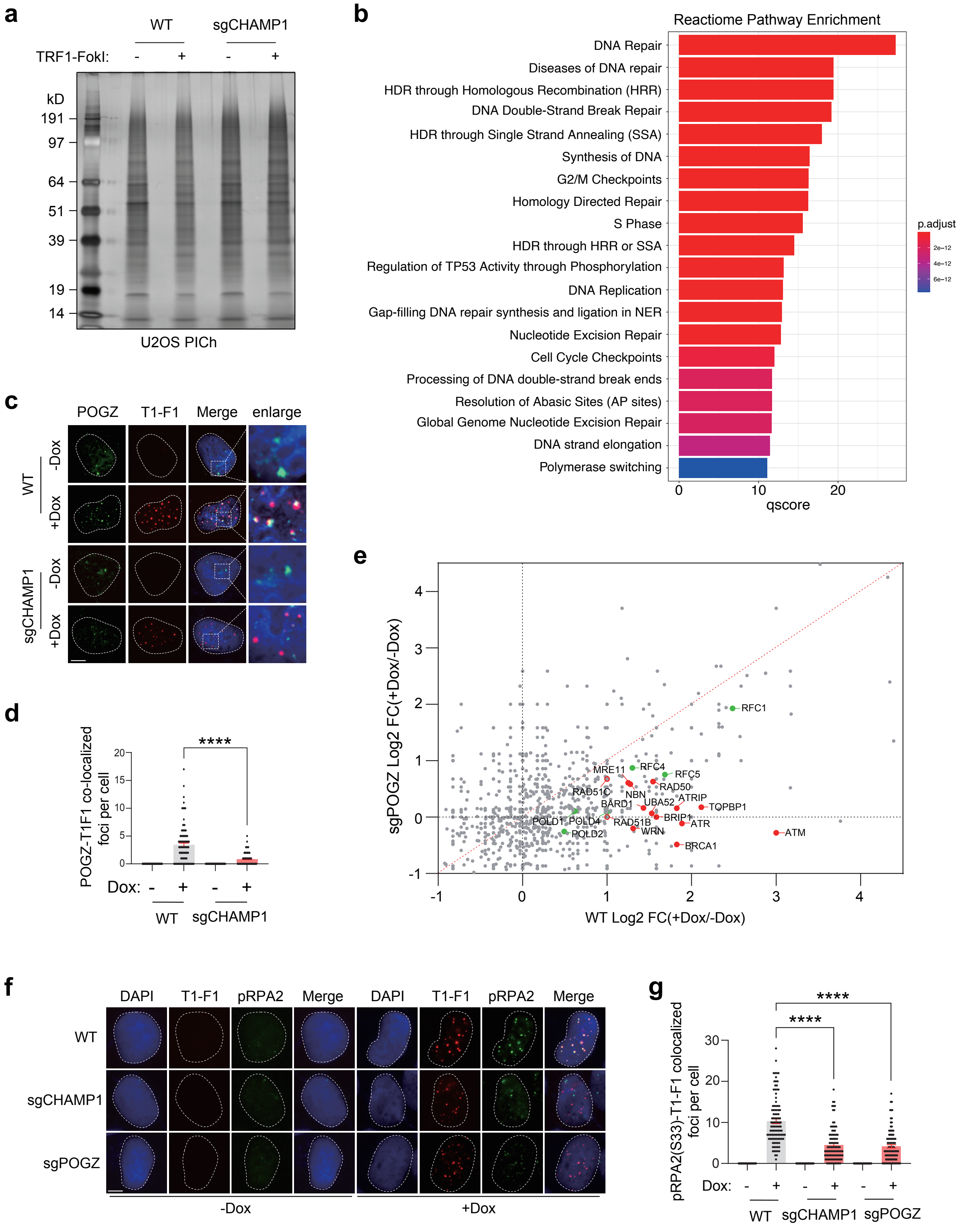

### Extended Data Fig. 5

**a**

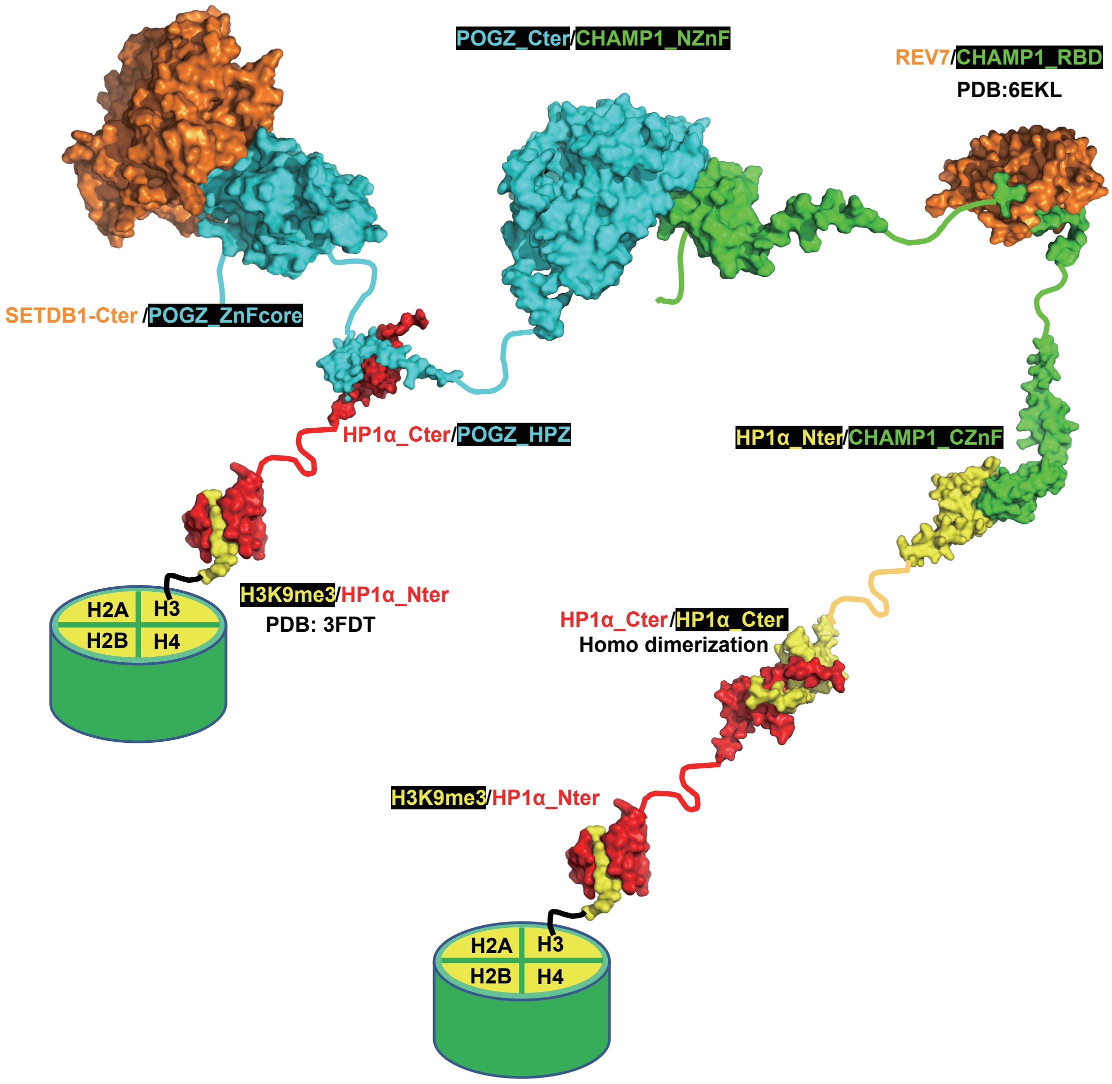


**b**

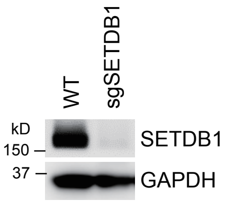


**c**

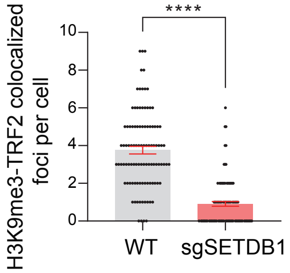

### Extended Data Fig. 6

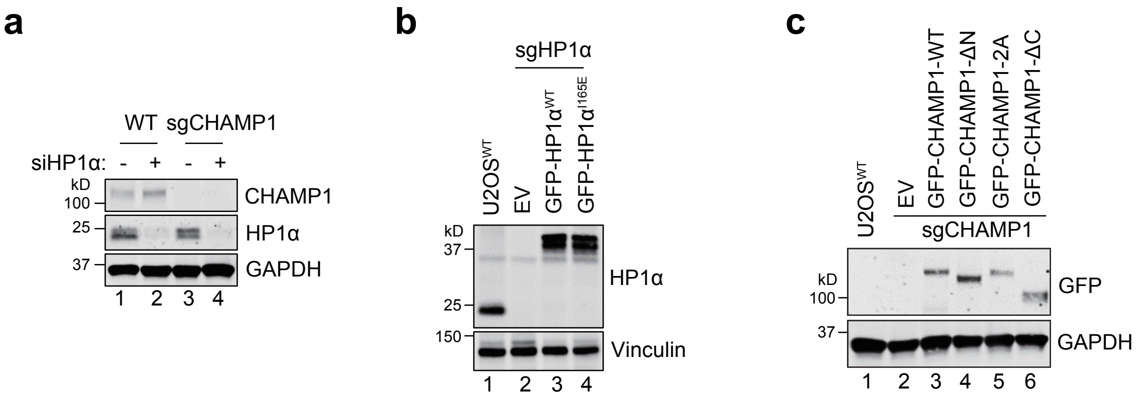

**Extended Data Fig. 7**

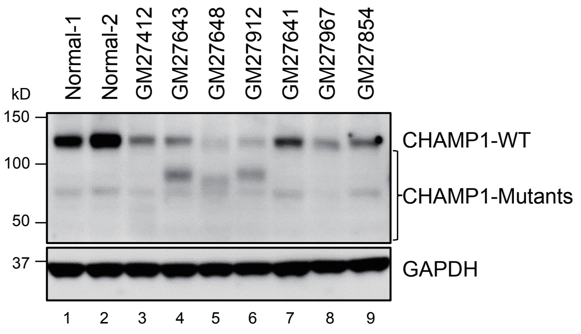
