## Supplementary material for "CHAMP1 complex directs heterochromatin assembly and promotes homology-directed DNA repair": Material and Methods: Material and Methods.docx

**Cell lines**

U2OS and U2OS TRF1-FOKI cell lines were grown in DMEM/F12 (Thermo Fisher) with 10% FBS (sigma) and 1% penicillin-streptomycin (Thermo Fisher). 293T, NIH-3T3 and HeLa S3 cell lines was grown in DMEM (Thermo Fisher) with 10% FBS (sigma) and 1% penicillin-streptomycin (Thermo Fisher). CHAMP1 patient lymphocyte cell lines from Coriell Institute were grown in RPMI1640 with 10% FBS (sigma) and 1% penicillin-streptomycin (Thermo Fisher). Cell lines were maintained in an incubator at 37 °C and 5% CO2 according to standard protocols. Cell lines were validated to be negative for mycoplasma contamination using the MycoAlert Plus Mycoplasma Detection Kit (Lonza) and the rapid MycoBlue Mycoplasma Detector (Vazyme).

**RNA interference, gene over-expression and CRISPR-mediated gene deletions**

DNA transfections were performed using Lipofectamine LTX (Invitrogen), while siRNA knockdowns were executed using RNAiMax (Invitrogen), following the manufacturer's guidelines. For single gene knockout, sgRNAs targeting candidate indicate genes were either be cloned into the pSpCas9 BB-2A-GFP (PX458) vector (GenScript) or introduced into the cell together with the Cas9 protein via electroporation (Lonza) according to the manufacturer’s protocol. After Cas9-gRNA PX458 plasmid transfection, GFP-positive cells were sorted using a BD FACSAria II cell sorter 48 hours post-transfection. GFP positive pool or single cells were screened for knockouts by western blotting.

**Cellular Fractionation and Immunoblot Analysis**

The cells were lysed using RIPA buffer supplemented with a cocktail of phosphatase and protease inhibitors from Roche. Cell lysates were separated by electrophoresis using NuPAGE 4-12% Bis-Tris gels (Invitrogen) and transferred onto nitrocellulose membranes. These membranes were then blocked with 5% BSA in TBST (Tris-buffered saline with Tween) and subsequently incubated with primary and secondary antibodies. The detection was accomplished using either chemiluminescence or fluorescence (LI-COR Biosciences).

For chromatin extraction, chromatin-bound extracts were obtained using a subcellular protein fractionation kit from Thermo. Band intensities were quantified using ImageJ.

**Immunofluorescence and IF-FISH Assays**

Cells were seeded onto glass coverslips placed in 24-well plates. Subsequently, they were either left untreated or exposed to 5Gy of ionizing radiation (IR). Following 6 hours, the cells were collected by pre-extraction with 0.5% Triton X-100 for 5 minutes, followed by fixation with 4% paraformaldehyde for 10 minutes at 4°C. After three PBS washes, a blocking step was carried out using 3% BSA in PBS for 1 hour at room temperature. This was followed by consecutive incubations with primary and secondary antibodies, conducted overnight at 4°C and 1 hour at room temperature, respectively. For IF-FISH assay, coverslips were first stained with the primary and secondary antibodies, fixed for 10min at room temperature and dehydrated in ethanol series. After denaturation at 85°C for 5min, coverslips were incubated with TelG-Cy3 or TelC-Alexa488 PNA probe (PNAbio) overnight at 37°C, then washed. Finally, the coverslips were mounted with DAPI (Vector Laboratories) and imaged using a Zeiss AX10 fluorescence microscope and Zen software. Foci were then counted, with a minimum of 100 cells assessed for each sample.

Telomere foci size and clustering was measured by ImageJ using a consistent threshold to images followed by binarization as described previously^1^. The sizes of the foci were quantified in square pixels for each telomeric focus within a nucleus, and the average size was computed for each analyzed nucleus.

**Clonogenic assay**

To evaluate the sensitivity of CHAMP1 or POGZ knockout cells to ATR or FANCM inhibitors, clonogenic assays were performed as described previously ^2^. Briefly, cells were seeded at 500 cells/well in 6-well plates. Drugs at the shown doses were added after 24 hours and cells were permitted to grow for 8-10 days. Colony formation was scored by fixing and staining with 0.5% (w/v) crystal violet in 20% methanol.

**CellTiter-Glo assay**

To evaluate the sensitivity of CHAMP1 patient lymphocyte cells to PARP inhibitors (PARPi) and ionizing radiation (IR), the short-term CellTiter-Glo survival assays were performed as described previously ^2^. For IR sensitivity, cells were exposed to indicated dose of IR and then plated in 96-well plates at a density of 800 cells per well. For PARPi sensitivity, cells were initially seeded in 96-well plates at a density of 800 cells per well, and then were treated with drugs at the indicated concentrations after 12 hours. Three days later, cellular viability was assessed using CellTiter-Glo (Promega). Survival at each dose of IR or drug concentration was calculated as a percentage relative to the corresponding untreated control.

**Immunoprecipitation**

After transfection for 48 hours, 293T cells were subsequently collected and subjected to lysis using NETN lysis buffer containing a proteinase and phosphatase inhibitor cocktail (Thermo, 1:100) for 30 minutes on ice. Following this, the lysed samples were incubated overnight at 4°C with an antibody-bead conjugate, which consisted of GFP-Trap_A (Chromotek). The beads were then thoroughly washed four times with NETN buffer, and the immunoprecipitated materials were eluted by boiling. Western blot analysis was conducted to detect the immunoprecipitates, and the intensities of the resulting bands were quantified using ImageJ.

**Chromatin immunoprecipitation (ChIP) and qPCR analysis**

ChIP assays were performed, as previously described. Initially, cells were treated with 1% formaldehyde for 10 minutes at room temperature to cross-link proteins to DNA, and the reaction was then halted by adding glycine to a final concentration of 0.125 M for 5 minutes. For ChIP assay, the SimpleChIP Kits (Cell Signaling Technology # 9003S) were used. 5μg H3 and H3K9me3 antibody were used for each ChIP reaction. Purified ChIP DNA was used as template for qPCR using the primers corresponding to telomeric repeats. qPCR experiments were performed using the QuantStudio™ 7 Flex Real-Time PCR System (ABI) with 96-well PCR plates and sealing films (Vazyme).

**Telomeric DNA synthesis in G2 phase**

To visualize telomeric DNA synthesis, U2OS cells were synchronized in G2 by treatment with 15 μM CDK1i (RO-3306) for 16h with or without the addition of Doxycycline. Cells were incubated with 20 µM EdU for an additional 2 hours, with or without 4-OH Tamoxifen, and then fixed with 4% PFA for 10 min at RT. Cells were permeabilized with 0.2% Triton X-100 in PBS for 10 min at RT, blocked with 10% normal goat serum for 1 h at 4°C then incubated overnight with anti-TRF2 antibody. Cells were then washed with 0.1% Triton X-100 in PBS, incubated with fluorescently-labeled secondary antibody and EdU was labeled with fluorescent dye using Click-iT EdU kit (Invitrogen) according to the manufacturer’s protocol. DNA with stained with DAPI and z-stack images were acquired using Zeiss AxioObserver microscope at 63x magnification. Images were analyzed using ImageJ software and the number of EdU+ TRF2 foci was assessed in at least 150 cells from 3 independent experiments.

**Proteomics of isolated chromatin segments (PICh)**

Telomere associated proteins were captured as previously described using PICh protocol^3-5^ with the following modifications. U2OS (~ 10^9^ cells) TRF1-FokI was induced with Doxycycline (40ng ml^-1^, Sigma-Aldrich, D9891) for 18hrs, then 4-Hydroxytamoxifen (1 µM, Sigma-Aldrich, H7904) was added for another 2hrs. Cells were fixed with 4% formaldehyde (Sigma-Aldrich) for 45 min with shaking at room temperature, washed three times with ice-cold 1× PBS, and scrapped down and collected in 1× PBS+ 0.05% Tween 20. The pellets were washed and dounced in sucrose solution. Cell pellets were resuspended in Triton solution and treated with RNase A overnight on a rotator at 4 °C, then washed with ice-cold 1× PBS for three times and with lysis buffer for twice. Then cell pellets resuspended in lysis buffer containing 1 mM PMSF and sonicated with BRANSON Digital Sonifier. Soluble fraction was collected and precleared with 1 mL Streptavidin Agarose (EMD Millipore, 69203-3) and then passed through Sephacryl S-400 HR column (GE Healthcare Life Sciences, 17060901). 0.01% of the sample was saved as “Input” for Western-blot. Soluble fractions were hybridized with 1500 pmol desthiobiotin labeled 2’-Fluor-RNA telomeric probes in a thermocycler. Hybridized chromatin was pooled and incubated with 900 µL Dynabeads™ MyOne™ Streptavidin C1 (Thermo Fisher, 65002) overnight at room temperature. Then bound chromatin was immobilized on a magnetic stand, 0.01% was taken as the “Unbound” fraction for Western-blot, followed by washing five times with normal salt lysis buffer at room temperature and one wash with low salt buffer at 42 °C. The telomere and associated proteins on beads were eluted twice with 450 µL elution buffer. The eluted samples, as well as the “Input” and “Unbound” fractions, were then precipitated, decrosslinked and boiled with NuPAGE LDS Sample Buffer (4×, Invitrogen). The samples were then analyzed by to silver staining, western-blot, and mass spectrometry (Taplin Biological Mass Spectrometry Facility at Harvard Medical School).

**AlphaFold2-Multimer structure prediction**

Protein sequences for CHAMP1 (accession# Q96JM3), POGZ (accession# Q7Z3K3), HP1α (accession# P45973) and SETDB1 (accession# Q15047) were retrieved from UniProtKB. Initially, full-length structural models of CHAMP1-POGZ, CHAMP1-HP1α, homodimer of HP1α and POGZ-SETDB1 complexes were predicted using a locally installed ColabFold (DOI: 10.1101/2021.10.04.463034, PMID:34265844, PMID:35637307) on a GPU machine. For each prediction, alphafold2 (AF2)_multimer_v3 with the default parameters for the run including num_recycles=20, num_models=5 and amber relaxation were used. After analyzing the resulting structural models of the full-length protein complexes for the most likely interface from each pair, a more focused prediction was performed using truncated protein sequences involving only protein domains that were part of the interface. For CHAMP1-POGZ complex structure prediction, CHAMP1 N-terminus Zn-finger domain (1-87 aa) and POGZ C-terminus domain (1021-1410 aa) sequences were used as input. For CHAMP1-HP1α complex structure prediction, CHAMP1 C-terminus Zn-finger domain (694-812 aa) and HP1α N-terminus domain (1-80 aa) sequences were used as input. For POGZ-SETDB1 complex structure prediction, POGZ Zn-finger core (468-693 aa) and SETDB1 sequence (560-1291 aa) that incorporates Methyl-CpG-binding domain (MBD) and two SET domains from the C-terminus were used as input. For HP1α homodimerization, HP1α C-terminus domain (109-180 aa) used as input sequence. For POGZ-HP1α complex prediction, we used POGZ HPZ-domain (791-850 aa) and HP1α C-terminus domain (109-180 aa) sequences were us as input. For each prediction, the resulting models were ranked based on the interface-predicted template modelling (iptm) score. Only the top models from each run were used for further analyses. All Structural analyses and figures were generated using Pymol (Schrödinger, Inc). All AF2 models were experimentally verified by immunoprecipitation protocol described above.

**QUANTIFICATION AND STATISTICAL ANALYSIS**

For the differential protein expression analysis of ALT childhood neuroblastoma tumors, we compared the ATRX-mutated ALT tumors with ATRX-wildtype ALT tumors using Limma^6^ (PMID: 25605792), utilizing log2-transformed LFQ protein intensity values. The LFQ protein intensity data for the ALT tumors, along with their ATRX mutation status, were acquired from Hartlieb et al.^7^ (PMID: 33627664). The volcano plot was then generated using ggplot2 in R.

All values are expressed as mean ± standard deviation (SD) or standard error of the mean (SEM) as indicated in figure legends. The statistical significance of differences was assessed by Student’s t-test for comparison of two groups, one-way analyses of variance (ANOVA) with Tukey’s test for comparison of multiple groups using Graphpad Prism 10 (Table S3).

1	Cho, N. W., Dilley, R. L., Lampson, M. A. & Greenberg, R. A. Interchromosomal homology searches drive directional ALT telomere movement and synapsis. *Cell* **159**, 108-121 (2014). <https://doi.org/10.1016/j.cell.2014.08.030>

2	Li, F. *et al.* CHK1 Inhibitor Blocks Phosphorylation of FAM122A and Promotes Replication Stress. *Mol Cell* **80**, 410-422 e416 (2020). <https://doi.org/10.1016/j.molcel.2020.10.008>

3	Dejardin, J. & Kingston, R. E. Purification of proteins associated with specific genomic Loci. *Cell* **136**, 175-186 (2009). <https://doi.org/10.1016/j.cell.2008.11.045>

4	Kan, S. L., Saksouk, N. & Dejardin, J. Proteome Characterization of a Chromatin Locus Using the Proteomics of Isolated Chromatin Segments Approach. *Methods Mol Biol* **1550**, 19-33 (2017). <https://doi.org/10.1007/978-1-4939-6747-6_3>

5	Zhang, T. *et al.* Break-induced replication orchestrates resection-dependent template switching. *Nature* (2023). <https://doi.org/10.1038/s41586-023-06177-3>

6	Ritchie, M. E. *et al.* limma powers differential expression analyses for RNA-sequencing and microarray studies. *Nucleic Acids Res* **43**, e47 (2015). <https://doi.org/10.1093/nar/gkv007>

7	Hartlieb, S. A. *et al.* Alternative lengthening of telomeres in childhood neuroblastoma from genome to proteome. *Nat Commun* **12**, 1269 (2021). <https://doi.org/10.1038/s41467-021-21247-8>
